# Optic Ataxia in Patients with Thalamic Lesions

**DOI:** 10.1101/2025.03.20.644311

**Authors:** Melanie Wilke, Hanna Eisenberg, Carsten Schmidt-Samoa, Shirin Mahdavi, Igor Kagan, Peter Dechent, Jan Liman, Hans-Otto-Karnath, Mathias Bähr

**Author notes:** **Corresponding Author:** Melanie Wilke, Ph.D., Department of Cognitive Neurology, Heart & Brain Center, University Medicine Goettingen, Robert-Koch-Str. 42, Goettingen, 37075, Germany. These authors contributed equally (Melanie Wilke, Hanna Eisenberg, Carsten Schmidt-Samoa).

## Abstract

Lesions in parietal cortex can strongly impair visually guided reach-grasping behaviour. A specific reaching deficit termed ‘Optic Ataxia’ (OA) occurs when eye and hand position are dissociated. Optic ataxia has typically been studied in patients with lesions in parietal cortex, neglecting potential thalamic contributions.

We here examined 28 acute stroke patients (age 58.9 ± 12.6 years) with circumscribed thalamic lesions for the presence of optic ataxia. We leveraged MRI-based lesion-symptom mapping to address the contributions of specific thalamic nuclei to visually-guided reaching deficits under foveal and peripheral viewing conditions.

Based on the cortical literature, we hypothesized that lesions in thalamic nuclei with strong connections to the inferior and superior parietal cortex, such as the ventrolateral nucleus and pulvinar might lead to optic ataxia.

In comparison with aged-matched healthy subjects (*n* = 40, age 60.6 ± 9.1 years), we identified five thalamic patients with optic ataxia, most pronounced for reaches with the contralesional hand into the contralesional space. While motor and grasping deficits and optic ataxia occurred frequently together, they did not always co-occur, and visual attention deficits could not account for the optic ataxia either. Comparing the lesion maps of patients with and without optic ataxia, the critical lesion site for optic ataxia was not restricted to one circumscribed thalamic region within the Morel atlas. Instead, it encompassed several medial and lateral nuclei within and around the internal medullary laminar (IML) complex. Interestingly, this region matches the so-called ‘central thalamus’, a functionally defined thalamic region that is considered a ‘higher-order’ nucleus complex. It receives afferent inputs from the cerebellum and brainstem regions and connects to fronto-parietal regions involved in eye movement control.

Taken together, our results suggest the critical importance of thalamic nuclei for the spatial transformation from eye-into body-centered coordinates.

## Introduction

Optic ataxia (OA) is a specific type of visuomotor deficit related to the interaction with visual objects in foveal vs. extrafoveal (i.e. peripheral) space. It is a compound syndrome that consists of misreaching into the visual periphery, difficulties in grasping objects and online correction of hand movements ^1,2^. The defining OA feature, however, is the difficulty to reach-grasp objects in peripheral space, i.e. when the gaze direction is dissociated from the reach goal ^3^. In unilateral lesion cases, reach errors in OA are most pronounced when patients use their contralesional hand to reach into contralesional space. Isolated optic ataxia in humans is rare, but patient cases with intact visual fields and preserved ocular and limb motor control have been described ^4^. Generally, reach-grasp functions are thought to rely on a distributed network of cortical regions such as motor, premotor, supplementary motor, superior parietal and dorsal occipital cortex as well as the cerebellum ^5,6^. Among those regions, optic ataxia has been specifically associated with lesions in the superior parietal lobule (SPL), intraparietal sulcus and the junction between the inferior parietal lobule (IPL) and superior occipital cortex ^4,7^. Since SPL and IPL are anatomically and functionally positioned between visual cortex and somatomotor-related prefrontal regions, they are thought to integrate visual and somatic information important for the control of movements ^8^. While the search for the neural mechanisms behind visually guided reaches primarily focused on cortical regions, there are several thalamic nuclei with strong connections to SPL/IPL. These thalamic connections are evidenced by anatomical tracer and microstimulation studies in macaques ^8–10^ as well as by anatomical and functional connectivity studies in humans ^11,12^. Specifically, SPL and IPL share reciprocal connections with somatomotor-related thalamic nuclei such as the ventrolateral (VL), ventroposterolateral (VPL) and centrolateral/centromedial nuclei (CL/CM), which also receive input from cerebellum and basal ganglia ^13,14^. Clinically, lesions in those thalamic nuclei have been associated with motor symptoms such as ataxia, dystonia, tremor, and motor learning deficits ^15–18^. In addition, SPL/IPL have strong connections with non-classical motor nuclei such as the lateral posterior nucleus and the dorsal pulvinar ^8,9,19^, which are thought to enable the efficient information transfer across visual and fronto-parietal cortices ^20,21^.

So far, it is unclear whether isolated thalamic lesions lead to optic ataxia. Studies of visually guided reach-grasp behavior after thalamic lesions are exceedingly rare, and the human literature consists almost entirely of individual case studies. In one case study, OA signified by misreaches towards contralesional, peripheral space was reported in a patient with left-sided lesions entailing the midbrain, cerebellum and thalamus ^22^. Another case study in a patient with bilateral medial pulvinar lesions reported abnormal reach-grasp behavior. The reach-grasp deficits of this pulvinar patient differed from optic ataxia however, as reach errors were not alleviated when he was allowed to look at the target ^23^. Two studies in non-human primates reported ‘optic ataxia’ following pharmacological inactivation or ablation of the pulvinar based on persistent misreaches and grasping deficits^24,25^. Importantly, while imprecise reach-grasping is often considered part of the optic ataxia syndrome ^2^, the OA defining performance difference between foveal vs. peripheral reach was not tested in those monkey studies.

We here investigated whether thalamic lesions and -if so-which subnuclei lead to optic ataxia in humans. We tested 28 patients with circumscribed thalamic lesions and compared the lesion location in patients with and without optic ataxia. In contrast to the referred primate and human case studies, the current study investigated a reasonably sized group of thalamic lesion patients with stroke aetiology. In addition, we mapped the lesion locations on high-resolution structural MRI and the Morel atlas, which allows anatomical precision and overcomes the narrow focus on single nuclei such as the pulvinar that characterizes previous work. Based on the cortical literature, we hypothesized that lesions in thalamic nuclei with strong connections to SPL/IPL cortex and known participation in integrating visual and motor information such as VL, pulvinar and LP might lead to optic ataxia. To our knowledge, this is the first patient population study that investigates the involvement of specific thalamic nuclei to visuomotor transformations required for gaze-dissociated reach movements.

## Materials and methods

### Study sample

We examined 28 patients with focal ischemic (*n =* 26) or haemorrhagic (*n =* 2) thalamic lesions. The stroke lesion group included 21 males and 7 females, 14 patients had predominately left and 14 patients had predominately right thalamic lesions (**Table 1**). Of those, six patients had an additional thalamic lesion in the opposite hemisphere, but all had a clinically relevant larger lesion on one side, which we use as reference for ipsi-vs. contralesional assignments (**Supplementary Tables 1** and **2**). The most affected lesion territories were the medial, lateral and posterior thalamus (see section on lesion characteristics below). The mean age of the patients was 58.9 years (range: 24.8-80.5, SD = 12.6). Testing for optic ataxia was performed within 10 days after stroke onset (median: 4 days, range: 2-9 days). MRI data were acquired within one day of behavioral testing. Patients were recruited from acute stroke patients at the ward of the Department of Neurology at the University Medical Center Göttingen. Following initial assessment including the clinical MRI or CT scan, patients with diffuse lesions, visual field deficits assessed by confrontation test, low vigilance or aphasia that would have interfered with understanding the task instructions were excluded. General exclusion criteria for patients and healthy controls were psychiatric disorders, chronic substance abuse or inability to lie still in the MR-scanner. In the final population, 39.3% had somatosensory (including paresthesia) deficits in the upper (contralesional) limb. 50% had a mild contralesional central hemiparesis of the upper limb (arm sinking in the holding test). The finger-to-nose test (FNT) was dysmetric in 14.3% of the patients. Light aphasia was present in 10.7% (**Table 1**). Signs of neglect, as defined by pathological results in two neglect tests was present in 1 patient. Testing for spatial neglect was conducted with several paper and pencil tests of the German neglect battery ^26^: (i) line bisection, (ii) line cancellation, (iii) star cancellation, the Apples test ^27^ and a computerized Posner task ^28,29^ (**Supplementary Methods)**. We also recruited a group of 40 age-matched healthy participants (20 male/20 female). The mean age of the healthy subjects was 60.6 years (range: 40.4-81.9, *SD* = 9.1). A two-sided t-test confirmed that the age of patients and healthy controls did not differ [*t(*66) = 0.63, *P* =0.53].

**Table 1.**
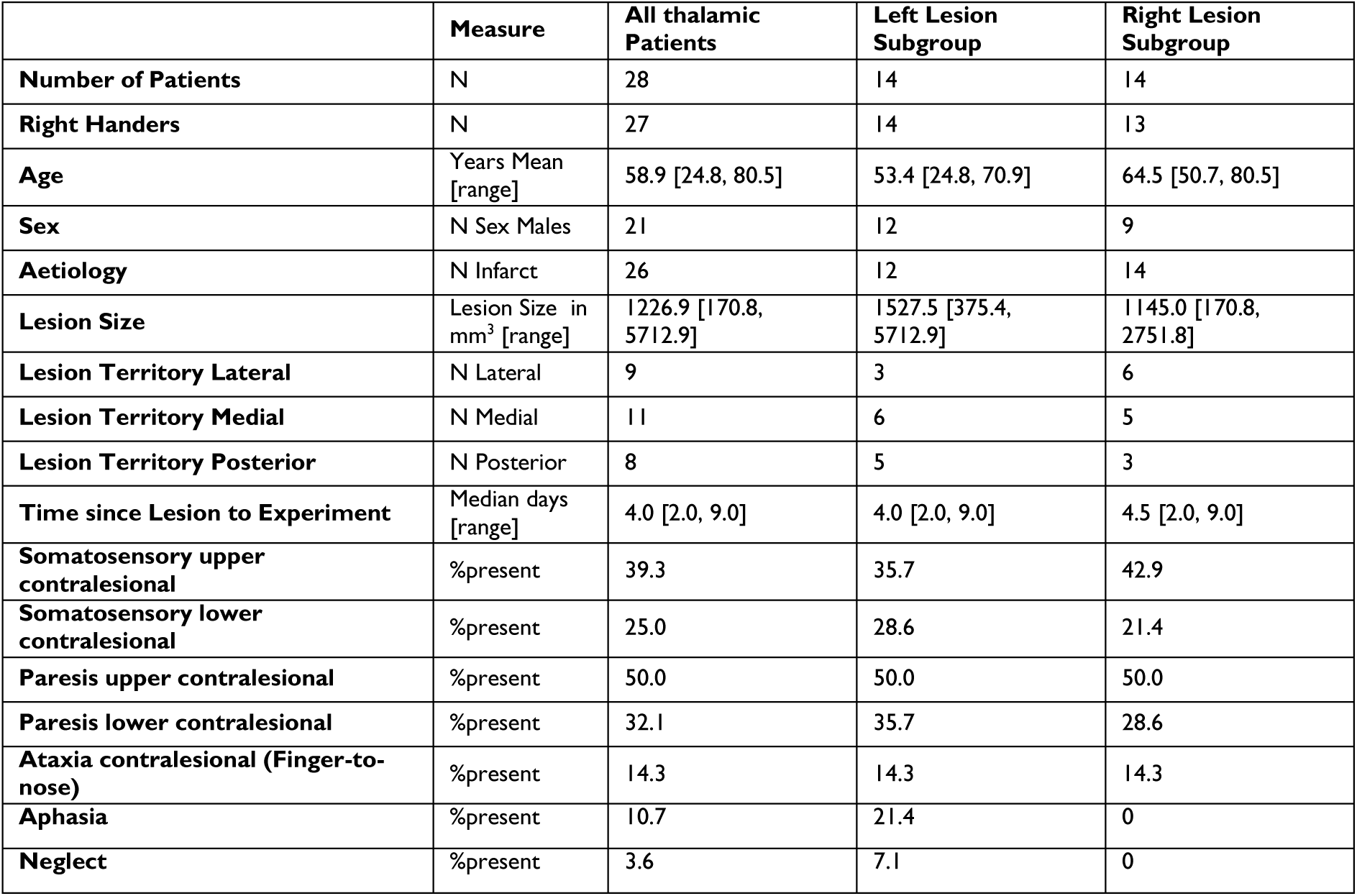
Demographic Information, lesion aetiology and clinical findings.

|  | Measure | All thalamic Patients | Left Lesion Subgroup | Right Lesion Subgroup |
| --- | --- | --- | --- | --- |
| **Number of Patients** | N | 28 | 14 | 14 |
| **Right Handers** | N | 27 | 14 | 13 |
| **Age** | Years Mean [range] | 58.9 [24.8, 80.5] | 53.4 [24.8, 70.9] | 64.5 [50.7, 80.5] |
| **Sex** | N Sex Males | 21 | 12 | 9 |
| **Aetiology** | N Infarct | 26 | 12 | 14 |
| **Lesion Size** | Lesion Size in mm<sup>3</sup> [range] | 1226.9 [170.8, 5712.9] | 1527.5 [375.4, 5712.9] | 1145.0 [170.8, 2751.8] |
| **Lesion Territory Lateral** | N Lateral | 9 | 3 | 6 |
| **Lesion Territory Medial** | N Medial | 11 | 6 | 5 |
| **Lesion Territory Posterior** | N Posterior | 8 | 5 | 3 |
| **Time since Lesion to Experiment** | Median days [range] | 4.0 [2.0, 9.0] | 4.0 [2.0, 9.0] | 4.5 [2.0, 9.0] |
| **Somatosensory upper contralesional** | %present | 39.3 | 35.7 | 42.9 |
| **Somatosensory lower contralesional** | %present | 25.0 | 28.6 | 21.4 |
| **Paresis upper contralesional** | %present | 50.0 | 50.0 | 50.0 |
| **Paresis lower contralesional** | %present | 32.1 | 35.7 | 28.6 |
| **Ataxia contralesional (Finger-to-nose)** | %present | 14.3 | 14.3 | 14.3 |
| **Aphasia** | %present | 10.7 | 21.4 | 0 |
| **Neglect** | %present | 3.6 | 7.1 | 0 |

Subjects’ written informed consent was obtained according to the Declaration of Helsinki and the study was approved by the medical ethics committee of the University Medical Center Goettingen, Germany. All patients signed an additional agreement to present their videos in scientific publications.

### Assessment of stroke symptoms of interest

Neurological assessments of somatosensory, motor and aphasia symptoms were performed by clinicians on the ward and were later derived from the clinical documentation. Somatosensory symptoms included all documented abnormalities of touch, pain and temperature sensation on either arm/hand (‘upper’) and legs (‘lower’) and reported dysesthesia/paraesthesia such as tingling and numbness. Motor evaluation included muscle tone and reflex status. Muscle strength was tested on arms (‘upper’) and legs (‘lower’) with the Medical Research Council (MRC) Scale for Muscle Strength and graded 0-5 (https://www.ukri.org/councils/mrc/facilities-and-resources/find-an-mrc-facility-or-resource/mrc-muscle-scale/). Ataxia refers to the dysmetria in the finger-to-nose tests. Presence of a visual field defect was assessed by finger perimetry. Grasping performance was tested in our laboratory with small objects and assessed by scoring the movies, similar to our previous pulvinar inactivation/lesion experiments in monkeys and humans ^23,30^ (**Supplementary Methods, Supplementary Fig. 1**).

### Optic Ataxia: main task description

The optic ataxia task was adopted from published optic ataxia guidelines ^3^. The task consisted of a pen being held into the peripheral visual field by the examiner standing behind the subject (**Fig. 1**), recorded by a second examiner using a digital camera. Participants were instructed by the experimenter on which hand to use. The pen was always presented in an upright orientation and peripheral with respect to the subject’s body with varying height between hand and elbow. Participants were either instructed to look at the pen as soon as they noticed it and then to reach for it (‘foveal reach’) or to keep fixating the camera while reaching for the pen (‘peripheral reach’). The examiner behind the camera controlled the performance and the trial was repeated if hand or eye instructions were not met. In addition, trials were labelled as valid or invalid during the video-based offline scoring based on this criterion. The median of valid trials across participants and for each hand/field and viewing condition ranged between 11-16 trials.

**Figure 1.**
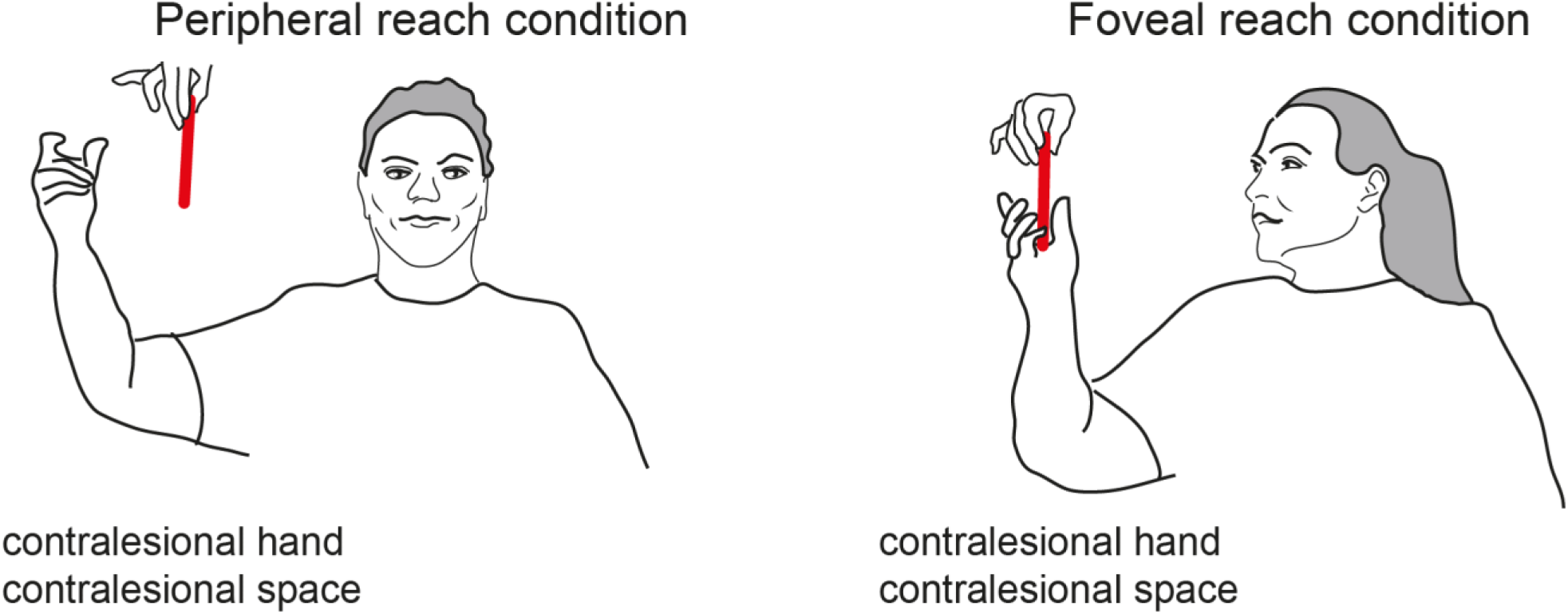
Optic Ataxia Task. One examiner presented a pen to the participant while the other experimenter stood behind the video camera (not shown) and observed the performance of the patient including the eye position. The pen was always presented in the peripheral space relative to the participant but varied in height. Shown is a sketch of video snapshot of an exemplary patient with optic ataxia from the current study (N063). The patient had a lesion in the left thalamus that encompassed CM, VPL and Pulvinar. The patient showed gross misreaches when the pen was presented in peripheral vision (eye fixation on the camera), while performing almost normal reaches under foveal vision (when allowed to orient eyes and head towards the object).

### Analysis of the Optic Ataxia task

#### Reach error scores

Individual error rates were analysed by two independent raters who scored each trial offline from the recorded videos. Error categories were (in raw error points given by the rater): fluent (0), slowed/insecure (1), corrected during (2) / after first (3) / second reach (4) / failed to reach (5) (category 5 never occurred). Reach-grasp errors and Optic Ataxia scores were calculated similar to Borchers et al. ^3^. First, a percentage mean error score was calculated for each of the eight (2×2×2, hand/space/gaze requirement) conditions using the formula mean error score = sum of trial error scores / (number of trials x 5) x 100. The error scores thus range from 0% (no error in any trial) to 100%. Higher values reflect worse reach performance. As some of the literature discriminated between corrected and uncorrected reaches ^4,7^, we also calculated the percentage of corrected and uncorrected reach errors. To this end, we categorized the errors into ‘fluent’ (score: 0), ‘corrected’ (slowed, insecure and corrected during reach, score: 1-2) and ‘uncorrected’ (scores 3-5), i.e. when the hand stopped at the wrong position and was corrected only in later reaches.

#### Optic Ataxia scores

In order to derive optic ataxia (OA) scores, a difference value was calculated for each hand/field combination by subtracting the individual mean error score in the foveal condition from the score in the peripheral condition. By the subtraction of viewing conditions to yield OA scores, primary motor factors such as paresis or tremor should cancel out ^3^. All scores were converted into ipsilesional and contralesional hand/space. For example, in a patient with a lesion in the left thalamus, the ipsilesional hand/space would be left and the contralesional hand/space right. In order to match the healthy controls (HCs), it was determined that in 50% of the HCs data from the left hand/space should be used for ipsi hand/space. The HC group assignment to ipsi/contra was based on 100000 permutations, which produced balanced mean and SD values for ipsi- and contralesional scores in the HCs.

### Structural brain imaging and lesion analysis

#### Anatomical MRI acquisition

MRI data were collected with a 3 Tesla MR system (Magnetom TIM Trio, Siemens Healthineers, Erlangen, Germany using a 32-channel phased-array head coil, or after scanner upgrade: Magnetom Prisma^fit^ using a 64-channel head coil). Three-dimensional (3D) anatomical datasets at 1 mm³ resolution were acquired in sagittal orientation with T1-weighting (turbo fast low angle shot (tFLASH), repetition time (TR): 2300 ms, inversion time (TI): 900 ms, echo time (TE): 2.96 ms (2.98 ms on Prisma^fit^), flip angle 9°) and with T2-weighting (fluid-attenuated inversion recovery (FLAIR), TR: 5000 ms, TI: 1800 ms, TE: 394 ms, integrated parallel acquisition technique: factor 2).

#### Anatomical lesion mapping

Lesions were manually segmented on FLAIR images using MRIcron. FSL 5.0.7 was used to coregister individual FLAIR images linearly to corresponding T1-weighted (T1w) images (FLIRT). T1w images were normalized to the MNI152 template (brain extraction (BET), 12-parameter affine (FLIRT) and non-linear registration (FNIRT), with lesions masked during registration). The transformation matrices were applied to the whole head T1w and FLAIR images and the lesion masks. Finally, the resulting images were upsampled to 0.5 mm isotropic resolution. A digitized version of the Morel atlas of the thalamus,^31,32^ registered to the 0.5 mm MNI152 template, allowed for visualization and analysis of the thalamic lesions with respect to thalamic nuclei (**Supplementary Methods**). For analysis in ipsi-/contralesional space, left thalamic lesions were mirrored along the midsagittal plane in MNI152 space. Individual scans and results of the lesion mapping are shown in **Supplementary Fig. 2 and Supplementary Table 1.**Statistical Analyses

#### Demographic, clinical and behavioural statistical data analysis

In order to assess whether optic ataxia occurs in thalamic patients at the population level and to derive pathological scores, we performed group comparisons between age matched-healthy controls, thalamic stroke patients and the subgroups of these patients with and without optic ataxia. In order to test whether the reach errors and optic ataxia scores are normally distributed, a Shapiro-Wilk test was performed and showed that the distribution of those variables departed significantly from normality (all *P-*values < 0.05 in both, HC (*df* = 40) and patients (*df* = 28)). Thus, group comparisons were performed using the non-parametric Mann-Whitney U-test.

#### Lesion characteristics and clinical profile of the patients

Thalamic lesions centered in 9 patients on the medial thalamus, in 11 on the lateral and in 8 on the posterior thalamus (**Supplementary Tables 1-2**). From the medial group, which includes the intralaminar nuclei (ILN), the centromedian nucleus (CM) (23 patients, 82.1%), central lateral (CL) (14 patients, 50%), parafascicular nucleus (Pf) (13 patients, 46.4%) and the mediodorsal nucleus (MD) (11 patients, 39.3%) were the most frequently affected. From the lateral portion, the ventral posterior (VP) (20 patients, 71%), ventral lateral (VL) (15 patients, 53%) and the ventral medial (VM) nucleus were damaged in 11 patients (39%). From the posterior group, the pulvinar and the lateral posterior nucleus were damaged in 12 (43%) and 11 patients (39%), respectively. The majority of patients had unilateral lesions, while 6 patients also had some damage in the other hemisphere. **Fig. 2** shows the overall lesion pattern for the entire group.

**Figure 2.**
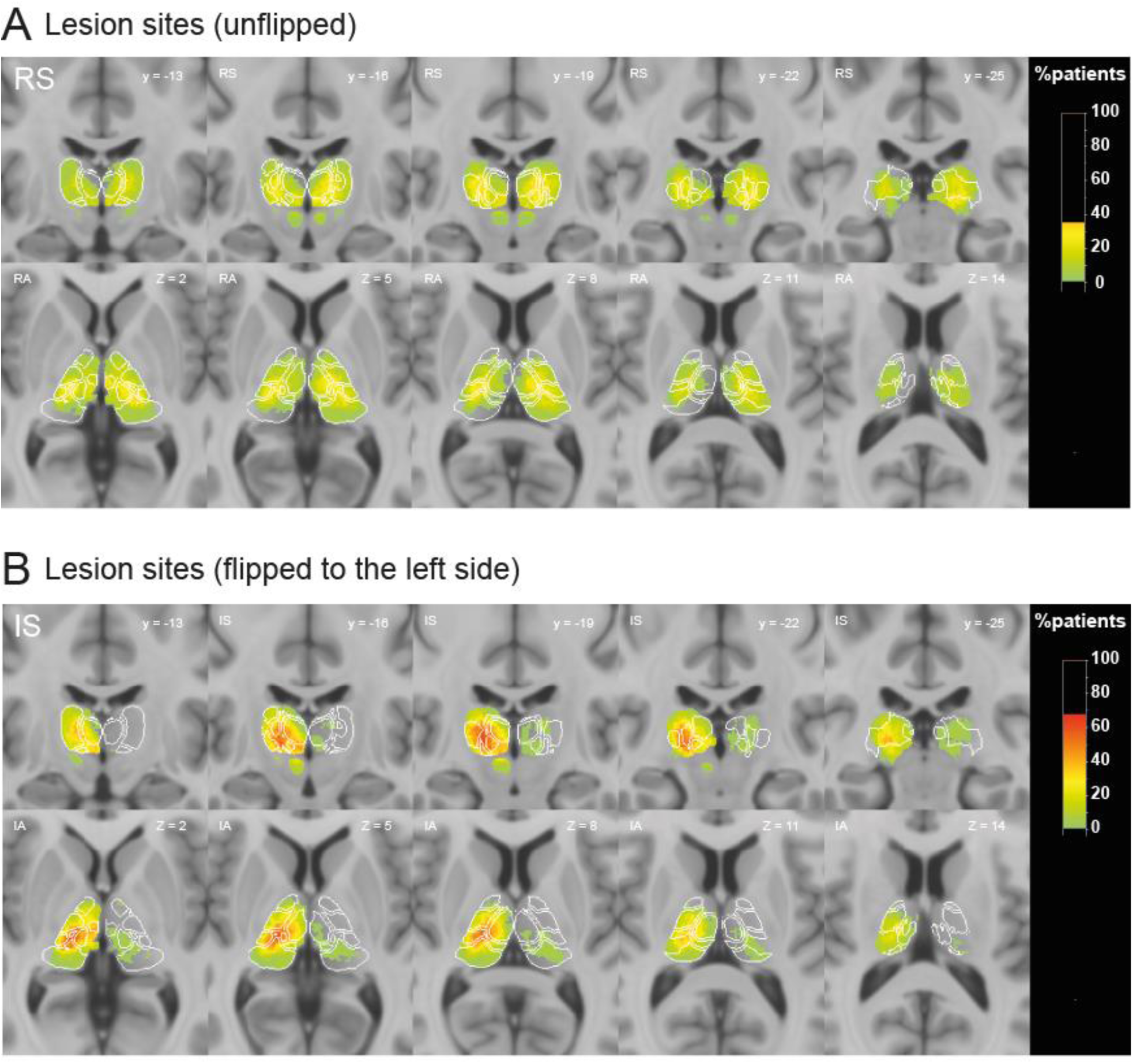
Lesion frequency map of the thalamic lesions in the whole group of stroke patients (*n =* 28) **(A)** Overlay lesion maps as they appear in the left and right hemisphere (radiological convention). **(B)** Lesions flipped so that the predominant lesion always appears in the right hemisphere. Upper panels in (A) and (B) show the coronal, the lower panels the axial slices. Coronal slices from anterior to posterior (left to right). Axial slices from top to bottom. Lesions were manually segmented on FLAIR MR images using MRIcron and transformed into MNI space. The colorbar specifies the percentage of patients with lesions in each voxel, with green indicating the fewest, and the hot colours (orange-red) a higher number of patients.

## Results

We examined 28 patients with circumscribed thalamic lesions. Participants were asked to reach and grasp a pen presented at different peripheral locations (**Figure 1**). To test for optic ataxia, trials that required eye fixation on the camera in front of the patients, and trials in which patients were asked to look at the pen, were interleaved. Reach performance in each patient was assessed by independent raters scoring the videos for reach smoothness and precision. The reach results are shown in **Fig. 3**.

**Figure 3.**
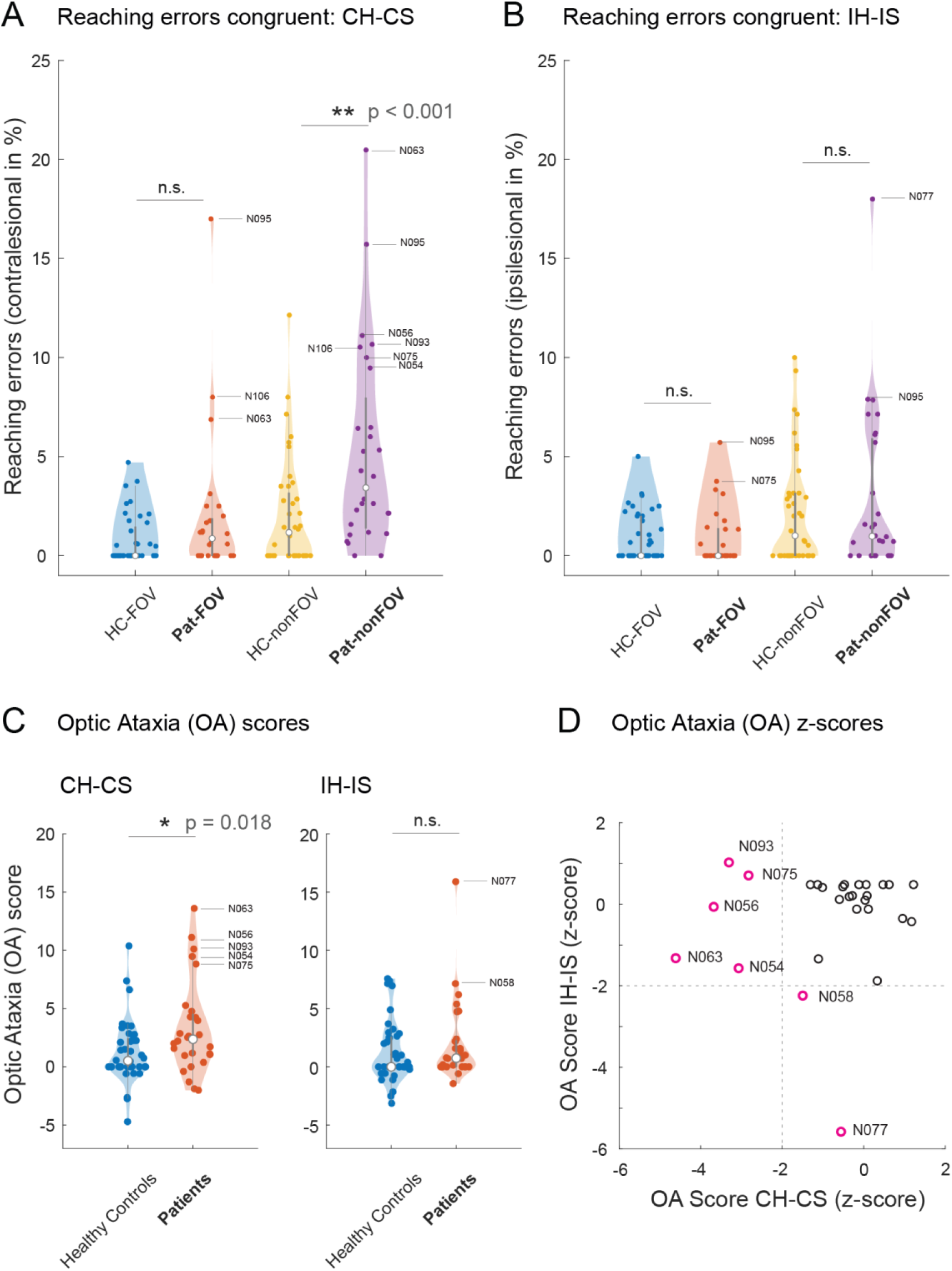
Reaching errors and optic ataxia scores as a function of hand and space and viewing conditions. Congruent condition refers to the same hand and space **(A)** Reaching errors for the contralesional hand and space (CH-CS), comparing foveal and non-foveal conditions. **(B)** Reaching errors correspond to the ipsilesional hand and space (IH-IS). In (A) and (B), ‘-FOV’ refers to the foveal condition, where subjects were allowed to look at the pole. ‘-nonFOV’ refers to the non-foveal (peripheral) condition where subjects fixated on the camera. Note the higher error rates in the non-foveal condition in both patients and controls. Importantly, the patient group showed the largest numbers of errors for peripheral reaches with the contralesional hand and contralesional space. **(C)** Distribution of optic ataxia scores in patients and healthy controls as a function of hand and viewing condition (CH-CS (left panel) and IH-IS (right panel)). Violin plots show the distribution of optic ataxia scores in patients and healthy controls. Optic ataxia scores represent the difference of reach errors in the peripheral (non-foveal) condition minus reach errors in the foveal condition, calculated separately for each hand/space combination. Note the shifted median error scores for this contralesional condition and the few patients with higher error scores than the main group. Gray bars represent the 95% confidence interval. **(D)** Scatter plots showing the z-scored contra- and ipsilesional optic ataxia scores in the patient group. As in neuropsychological convention, the sign of the z-scores was flipped so that negative values represent worse performance. In all plots (**3A-D**), each dot represents as single patient, the labelled ones performed worse than <-2 SD from the healthy control group, which were also significant in the Crawford-Garthwaite test for single subjects ^33^. In the violin plots **3A-C**, the white dots represents the median, the thick grey bar the interquartile range, the thin grey lines (‘whiskers’) represent the 1.5 times the IQR above the third quartile and below the first quartile. The shape of the violin depicts the density of scores in each group (broad means higher frequency).

### Reaching errors in foveal and peripheral conditions

We first compared the reach performance between patients and healthy controls. In the optic ataxia literature, reach errors are typically distinguished by hand, space and viewing condition ^3,4^. From the video ratings, an individual percentage error score was calculated for each of the hand, space and viewing conditions (**Materials and methods**). For simplicity we focus the main figures on the OA-relevant congruent hand-field conditions, where hand and side of the pen match ^3^. Those two critical conditions were also used for Bonferroni corrections for multiple comparisons. Medians and ranges for all groups including the crossed (incongruent) conditions are presented in **Supplementary Table 3**.

#### Reaching errors: Foveal condition

Apart from a few exceptions, healthy subjects rarely made any reach errors when they were allowed to look at the pen, yielding median error scores of 0 (assigned ‘contralesional’ hand/space combination [CH_CS]: range 0-4.7%; assigned ‘ipsilesional’ hand/space combination [IH_IS]: 0-5.0%) (**Fig. 3A, Supplementary Table 3**). Similarly, only few thalamic patients made errors in the foveal condition, either in the contralesional hand/space (CH-CS) (median: 0.9%, range: 0-17%) or ipsilesional hand/space combination (IH-IS) (median: 0.9%, range: 0-5.7%). Accordingly, the Mann-Whitney U test did not yield a significant result for the comparison between patients and the healthy control group in the foveal condition (CH-CS: *U* = 683.0, *P* = 0.099; IH-IS: *U* = 521.0, *P* = 0.593). In the foveal condition, the U-Test also lacked significance for the incongruent conditions (CH-IS: *U* = 618, *P* = 0.422; IH-CS: *U* = 521, *P* = 0.478). Separating the percentage of corrected and uncorrected errors yielded a similar picture. Very few errors occurred in the foveal condition for either hand/space combination.

#### Reaching errors: Non-Foveal (peripheral) condition

From the (cortical) optic ataxia literature, we expected the highest error rates for reaches with the contralesional hand to the contralesional space and for the peripheral viewing condition ^2,7^. Since reach-grasp movements are more difficult to execute when we don’t look at the objects, healthy subjects and patients had higher error rates in this peripheral (non-foveal) condition (**Fig. 3A, B; Supplementary Table 3**). Critically, as a group, patients made significantly more errors than the healthy controls when using their contralesional hand towards the contralesional space in this peripheral condition (median: 3.4%, range: 0-20.5%; *U* = 820.0, *P* < 0.001; **Fig. 3A**). None of the other hand-field combinations yielded a significant group effect (all *P* > 0.15). In terms of percentage of corrected errors in the non-foveal CH-CS condition, the median of the patients was 10.0% with a large range from 0 to 69%. Uncorrected errors were rare, resulting in an overall median of 0 (range: 0 to 7.1%) even in the CH-CS condition. In the ipsilesional hand/space condition, there were less corrected and uncorrected errors overall, yielding a median of 3.7% (range: 0-39.3%) of corrected and 0% uncorrected errors (range: 0 to 13.3%) (**Supplementary Table 3**).

### Optic Ataxia scores

The higher amount of reach errors in the contralesional peripheral condition already hinted at the occurrence of optic ataxia in the thalamic patients. The defining feature of optic ataxia is the disproportionate occurrence of errors when objects are in peripheral space, i.e. are not foveated ^2^. To test directly for optic ataxia, we next subtracted the individual error scores in the foveal condition from the error scores in the peripheral condition ^3^. As a group, thalamic patients had significantly higher optic ataxia scores than healthy controls when using their contralesional hand to reach into the contralesional space (*U* = 749.5, *P* = 0.018) (**Fig. 3C**). It is also clear from this figure that the significance at the group level is driven by five individual patients with high OA scores. Arguably, those patients are more important than the group mean and will be detailed below. On the group level, there was no significant deficit when patients used their ipsilesional hand and reached to the ipsilesional space (OA Score IH-IS, *U* = 639.0, *P* = 0.32). **Fig. 3D** shows the z-score transformed regression plot between the Optic Ataxia (OA scores) for the combination contralesional hand-contralesional space (CH-CS) and ipsilesional hand/ipsilesional space (IH-IS) conditions. A negative z-score indicates higher optic ataxia scores, and a threshold of −2 indicates a performance worse than two standard deviations from the healthy controls. Five thalamic patients had a poorer performance in the CH-CS condition and two patients a poorer performance in the ipsilesional (IH-IS) condition (**Fig. 3D**). Median and ranges for all hand/eye conditions are depicted in **Supplementary Table 3**.

### Selective case analysis

Since OA studies in thalamic patients are exceedingly rare, we here present some details of the five individual OA cases to show the range of space and/or hand-specific effects and their putative relationship to the thalamic lesion sites (**Fig. 4**). **Fig. 4A, B** depict the MR-based lesion centers of each patient, with a descriptive thalamic nuclei scheme next to it. Exact lesion locations for each patient are shown in **Supplementary Table 1**. **Fig. 4C, D** depict the optic ataxia scores and the proportion of corrected/uncorrected errors of each patient, respectively. The combination of contralesional hand/space (CH/CS) was typically the worst condition, while hand and space could be affected to varying amounts. In the group of the patients with contralesional OA, the majority of patients made mostly corrected errors (**Fig. 4D**). Somatosensory impairments, including misperceptions in the upper limbs were not a prerequisite for OA. Furthermore, an abnormal performance in the arm holding test (mild paresis) was often present, while this modest sinking of the arm with closed eyes could also be attributed to proprioceptive deficits. For example, patient N063 with the most pronounced deficits, had almost symmetrical bilateral lesions in the medial (CM, CL) and posterior portion (Pulvinar, LP), while the lateral lesions (VL, VM, VP) were more pronounced in the left hemisphere. In the standard clinical testing, the right lesion remained clinically silent such as tingling of the arm and dysmetria in the finger-to-nose test occurred on the right side only (see **Patient Descriptions in the Supplementary material**). In the optic ataxia task, N063 made a large amount of corrected errors in the non-foveal condition with the right hand. In those trials, only 30.9% trials with this hand were rated effective and fluent, while the remaining trials were slowed and ineffective but were corrected in flight. The slowed and corrected reaches with the right hand in the right space led to a pathological optic ataxia score in this patient (z = −4.6). The ipsi hand/field OA condition was not impaired (OA z-score: −1.32) (**Fig. 4**, middle panel). She also made uncorrected errors, while those are likely underestimated in our cohort, as patients sometimes looked at the pen when they did not reach it, in which case the trial was invalidated. Grasping data of the whole population are shown in the **Supplementary Fig. 3**. Only two of the five OA patients showed grasp impairments when they were allowed to look at the small items. Thus, on the thalamic level, OA and grasping impairments can dissociate. The extended case studies of each patient shown can be found in the **Supplementary Materials** (‘**Single Patient Descriptions’**). The two patients with ipsilesional deficits showed a pattern that is hard to interpret (data not shown): the first patient (N058) had only a minor deficit for both, the CH-CS and the IH-IS condition (CH-CS: z = −1.4; IH-IS: −2.24). Patient N077 (no neglect, VEP’s intact) had a strong ipsilesional space effect, exhibiting the strongest deficits for the IH_IS and CH_IS conditions (z = −5.58 and z = −9.74).

**Figure 4.**
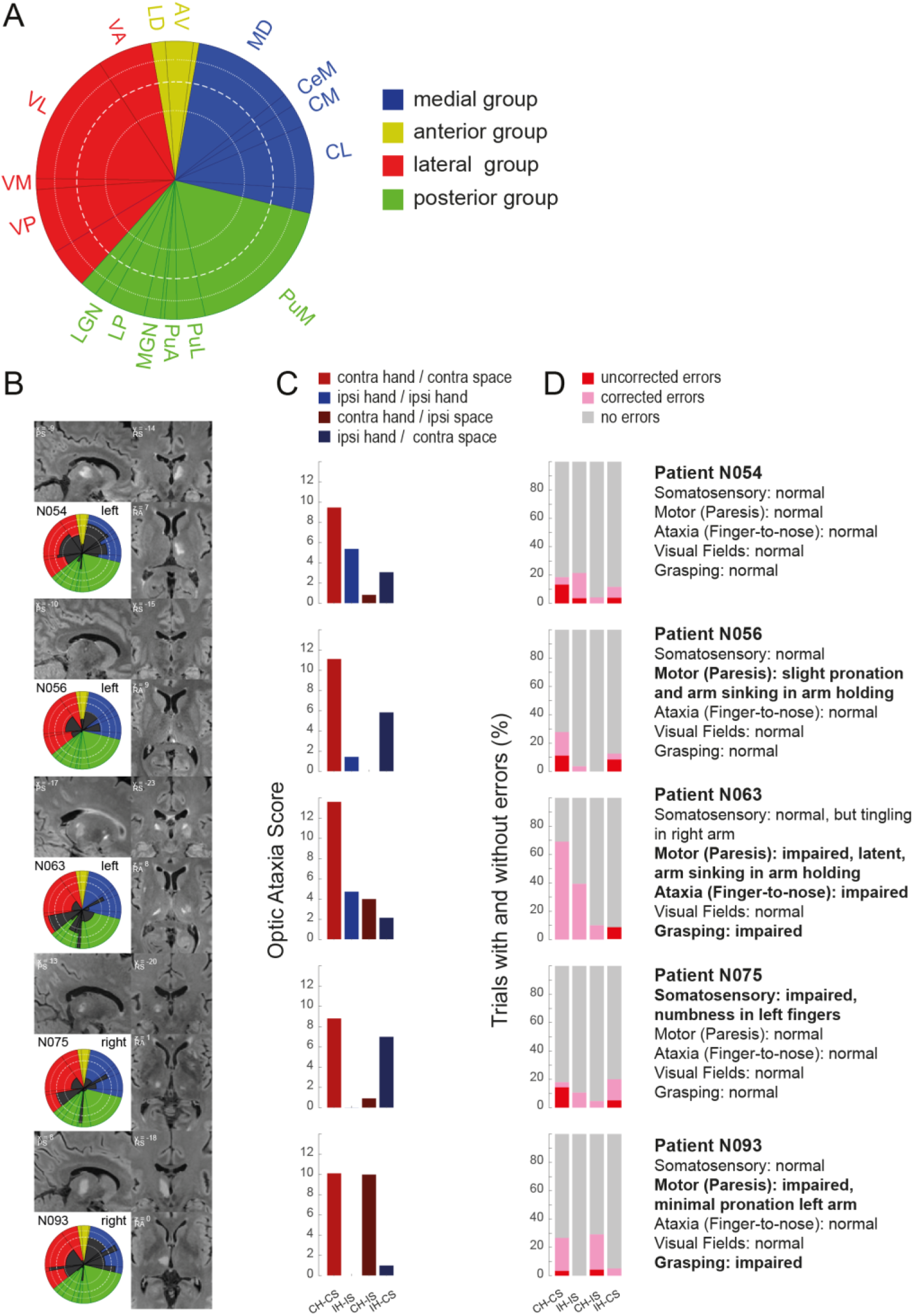
Selective Case Analysis. Patients with pathological Optic Ataxia scores in the CH/CS conditions. **(A)** Morel Atlas scheme. **(B)** Individual anatomical scans (FLAIR) were registered to the high-resolution MNI 152 template to visualize the thalamic lesions. Schema shows the distribution of the lesion on the morel scheme. **(C)** Optic Ataxia scores for each patient (non-foveal minus foveal errors), separated by hand and space. **(D)** Percentage corrected and uncorrected errors in the non-foveal condition. The text depicts the main clinical features of the patient. Abbreviations: CH-CS: contralesional hand-contralesional space), IH-IS: ipsilesional hand-ipsilesional space, CH-IS: contralesional hand-ipsilesional space, IH-CS: ipsilesional hand-contralesional space). The number of valid trials for each patient were as follows (Foveal/Non-Foveal): **N054**: CH-CS: 17/19, IH-IS:17/14, CH_IS: 13/12, IH_CS: 13/14; **N056:** CH-CS: 15/18, IH-IS: 20/15, CH_IS: 13/13, IH_CS: 15/13; **N063:** CH-CS: 16/21, IH-IS: 16/14, CH_IS: 14/10, IH_CS: 14/12; **N075**: CH-CS: 17/14, IH-IS: 16/19, CH_IS: 15/13, IH_CS: 15/10; **N093:** CH-CS: 20/15, IH-IS: 15/20, CH_IS: 13/12, IH_CS: 13/10.

### Patients with and without optic ataxia

We next focused on the question which clinical variables discriminate between thalamic patients with and without optic ataxia on the population level. To this end, we used the z-transformed optic ataxia scores based on the healthy control group. We then compared the patients who had an OA score worse than 2 standard deviations with the patients with normal z-scores (**Fig. 3C, D**). The group of patients with contralesional optic ataxia vs. without contralesional optic ataxia (OA, *n =* 5) did not differ in age, sex, etiology nor in their basic clinical parameters such as somatosensory, motor or grasping deficits (Mann-Whitney or χ2 Test, where appropriate, **Table 2**). Similarly, none of the comparisons reached significance when adding the two OA patients with ipsilesional OA and comparing the resulting OA group (*n =* 7) with the non-OA patients (*n =* 21) (all p > 0.1). Optic ataxia occurred after left or right thalamic lesions (**Table 1**), whether there is some laterality needs to be addressed with larger sample sizes. One contributing factor was whether the thalamic lesion was unilateral or bilateral. In the group of contralesional OA patients three out five (60%) had also a smaller lesion on the other side (**Fig. 4B**). In contrast, only 3 out of 23 (13%) patients without OA had a bilateral lesion (*chi-square* = 5.38, *P* = 0.02).

**Table 2.** Demographic and clinical data of all patients who presented with and without pathological optic ataxia scores.

|  | Statistical Measure | OA Patients | NonOA Patients | Statistical test | P-values |
| --- | --- | --- | --- | --- | --- |
| Number of Patients | N | 5 | 23 |  |  |
| Handedness | %percent right | 100 | 96% | chi-square ($\chi^2$) | 0.635 |
| Age | years | 55.8 [24.8, 71.2] | 59.6 [32.3, 80.5] | Mann-Whitney U | 0.88 |
| Sex | N Sex Males | 4 | 17 | chi-square ($\chi^2$) | 0.776 |
| Aetiology | N Infarct | 4 | 22 | chi-square ($\chi^2$) | 0.331 |
| Lesion Side | Lesion Side (Left) | 3 | 11 | chi-square ($\chi^2$) | 0.866 |
| Bilateral Lesion | N present (percent) | 3 out of 5 (60%) | 3 out of 23 (13%) | chi-square ($\chi^2$) | **0.02** |
| Lesion Size (in mm3) | Lesion Size | 1802.0 [781.3, 2085.8] | 1089.1 [170.8, 5712.9] | Mann-Whitney U | 0.139 |
| Time since Lesion | Median days | 6.0 [3.0, 9.0] | 4.0 [2.0, 9.0] | Mann-Whitney U | 0.792 |
| Somatosensory_upper_contra | %present | 20 | 43.5 | chi-square ($\chi^2$) | 0.33 |
| Somatosensory_lower_contra | %present | 0 | 30.4 | chi-square ($\chi^2$) | 0.154 |
| Paresis_upper_contra | %present | 60 | 47.8 | chi-square ($\chi^2$) | 0.622 |
| Paresis_lower_contra | %present | 20 | 34.8 | chi-square ($\chi^2$) | 0.521 |
| Ataxia Contra | %present | 20 | 13.0 | chi-square ($\chi^2$) | 0.687 |
| Aphasia | %present | 20 | 8.7 | chi-square ($\chi^2$) | 0.459 |
| Neglect | %present | 0 | 8.7 | chi-square ($\chi^2$) | 0.635 |

\*denotes the significant p-values

### Lesion locations in patients with and without optic ataxia

In order to identify the critical lesion sites for patients with and without optic ataxia we next contrasted the lesions of the five patients with (contralesional) OA to thalamus lesions in patients without OA. **Fig. 5A** shows the most common lesion sites in the thalamic patients with and without optic ataxia. Based on the (cortical) literature we hypothesized that OA should occur in patients with lesions in thalamic nuclei with strong connectivity to SPL/IPL - i.e. VPL, VL and LP/pulvinar. Since we did not find a hemispheric trend for optic ataxia scores (Mann-Whitney U-test, *P* = 0.87, **Table 2**), we flipped all lesions to the right hemisphere. In the group of the 5 optic ataxia patients, 100% had a lesion that involved the lateral thalamus and in 80% (*n =* 4) also the medial thalamus (**Fig. 5A**, left panel). The posterior group was only involved in one OA patient. From the medial group, MD was damaged in 40% (MDmc in 40% and MDpc in 60%). Within the medial group, where the intralaminar nuclei (ILM) are placed according to the Morel atlas, CL and CM were affected the most frequently (80%). From the lateral group, VPL was affected in 100% and VL in 80%. Within the posterior group, Pulvinar or LP was damaged in only one patient. In order to identify the thalamic nuclei that are typically affected in patients with OA, it is important to directly contrast the patients with and without optic ataxia. Given the differing vasculature-induced differences in the likelihood of stroke affection, the relative percentages between lesions in OA vs. non-OA lesion controls is important. For example, a CL lesion was present in 80% of the OA patients, but only in 43.4% patients without OA (**Fig. 5A**, middle panel). Similarly, a VL lesion was present in 80% of the OA patients but only in 47.8% of the patients without OA. VPL was affected in all OA patients, but only in 52% of the non-OA patients. In pulvinar and LP the pattern is reversed: only 20% of OA patients had pulvinar or LP lesions, but 43.4% of patients without OA had a lesion there **(Fig. 5A**, middle panel). Thus, to identify the thalamic nuclei that are commonly lesioned in OA patients but spared in the OA, we subtracted the lesion overlays of patients without OA from the OA patients (**Fig. 5A**, right panel, **Fig. 5B**). The non-flipped data on the original hemisphere are shown in **Supplementary Fig. 4**. The thalamic structures that were commonly damaged in patients with OA but were typically spared in patients without OA were MDl, CL, CM, VPL and VL, with a concentration at the border between them, i.e. the ILM. Pulvinar nuclei did not belong to this group of lesions that were commonly damaged in OA patients but were typically spared in patients without OA. Schematic borders of the thalamic nuclei can be found in **Supplementary Fig. 5**.

**Figure 5.**
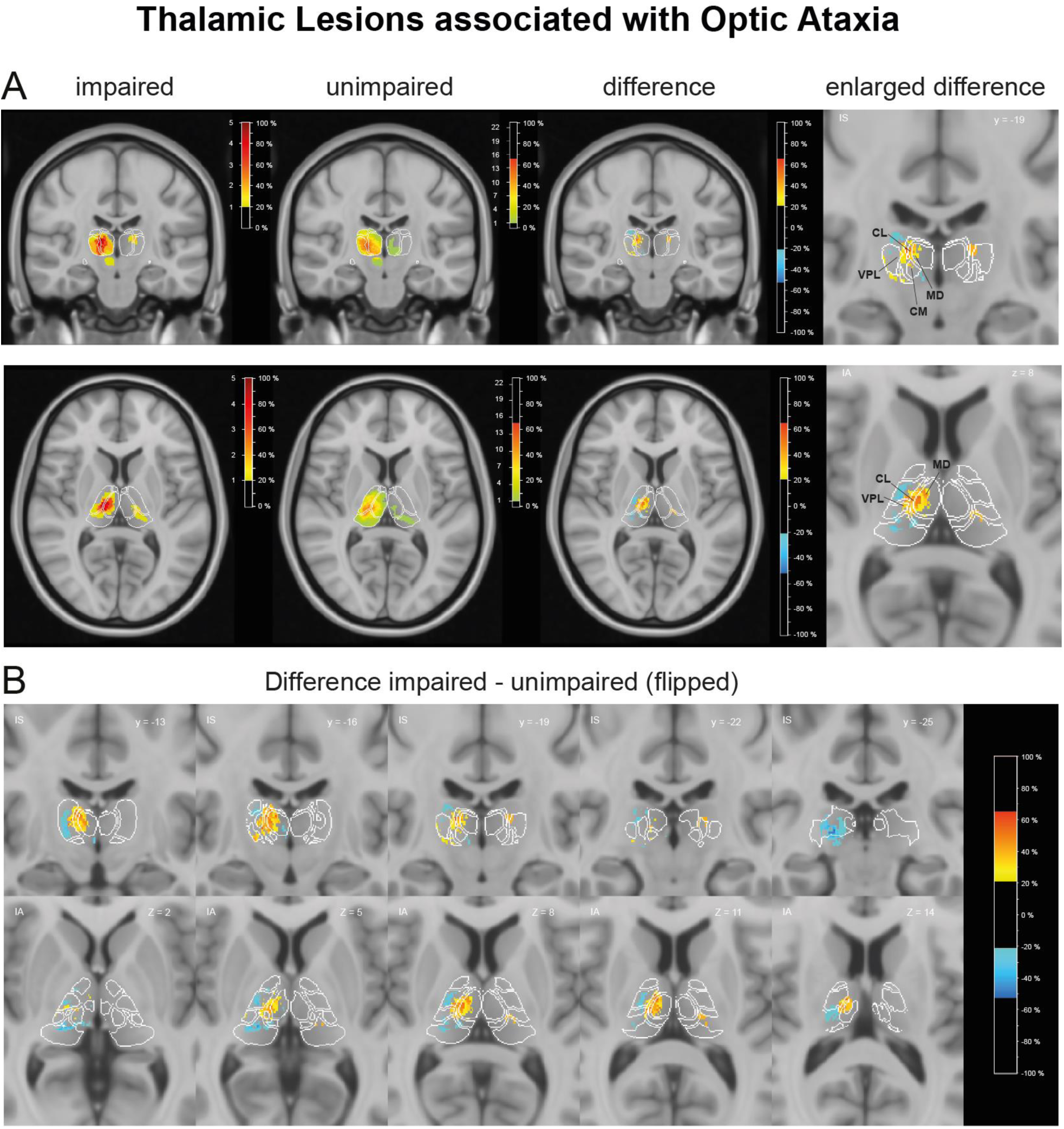
Lesion subtraction plots of the patients with and without optic ataxia. **(A)** Lesion overlap of the patients with and without optic ataxia. Primary lesions were flipped to the right hemisphere (see **Supplementary Fig. 4** for non-flipped maps). Shown are the impaired patients (left), unimpaired (middle) and the difference (right panel). The rightmost inset shows the enlarged difference of the slice. **(B)** Percentage of overlapping lesions of the optic ataxia patients after subtraction of the patients without optic ataxia. Coronal (upper panel) and axial (lower panel) slices are shown. The colorbar specifies the percentage of patients with lesions in each voxel, with blue-green indicating the fewest, and the hot colors (orange-red) a higher number of patients with an OA deficit. White outlines represent the borders of thalamic nuclei as defined in the digital version of the Morel-atlas normalized to the 0.5 mm MNI152 T1 template. Abbreviations: VL, Ventral lateral posterior nucleus; VPL, Ventral posterior lateral nucleus; MD, Mediodorsal nucleus; CL, Central lateral nucleus; CM, Centromedian nucleus. Thalamic nuclei on selected coronal and axial slices are depicted in **Supplementary Fig. 5**. Lesion locations for each patient and summary percentages are shown in **Supplementary Tables 1 and 2**.

### Control analyses

Although the lesion subtraction method should remove unspecific effects that are due to the general lesion distribution, we conducted several control analyses to address the specificity in our sample. To this end we computed the lesion subtraction for patients with and without paresis of the upper limb (*n =* 14 vs. *n =* 14). **Supplementary Fig. 6 A** and **B** shows that VL and VPL were the most commonly affected in patients with contralesional limb problems. In contrast, lesions of patients with and without contralesional somatosensory deficits (including paresthesia, pain etc. (*n =* 11 vs. *n =* 17)) centered on VPL and the anterior pulvinar. Similarly, when we computed the lesion subtraction to compare patients with and without deficits in the smarties grasping task (*n =* 7 vs. *n =* 21), sites that were commonly affected in patients with grasping difficulties but were typically spared in patients without also centered on VPL and anterior pulvinar (**Supplementary Fig. 7**. This suggests, the grasping deficits may be due to the somatosensory and/or sense of position impairments in those patients. Nucleus delineations for all panels are available in **Supplementary Fig. 5**. Thus, in contrast to the optic ataxia other deficits such as motor, somatosensory or grasping symptoms affected different thalamic regions and did not center on the border between lateral MD and VL. All difference maps were distinct from the difference we found relevant for optic ataxia.

## Discussion

Our analysis of 28 stroke patients revealed that optic ataxia occur as a result from circumscribed thalamic lesions. In fact, in 5 of these 28 thalamic patients we found typical OA behavior. Reaches with the contralesional hand towards the contralesional space were particularly impaired, similar to a number of previous studies in OA patients with cortical lesions ^1,7^. This combination of hand and field effects in (unilateral cortical lesion-induced) OA has been taken as evidence that OA is caused by a problem to transform visual input into motor output and/or between spatial coordinate frames ^6,34^. Optic ataxia in our thalamic patients could not be explained by either primary sensory (visual or somatosensory), motor deficits or spatial neglect. The direction of misreaches was not necessarily towards the central fixation, thus we did not observe systematic ‘magnetic misreaches’ ^35–37^. Although grasping deficits and OA occurred frequently together, they did not always co-occur; similar to some optic ataxia patients with cortical lesions^38^. The lesion subtraction analysis of 28 thalamic stroke patients with and without optic ataxia showed a region in both hemispheres that centered around the ILM and included parts of CL, lateral MD (i.e. MDpc), VPL and the medial portion of VL. Pulvinar lesions, despite its strong connectivity with parietal cortex, were surprisingly not associated with OA. The lesion pattern associated with OA was clearly distinct from the lesion patterns that led to paresis, somatosensory or grasping deficits with small items in the center of gaze. Instead, lesion sites associated with grasping deficits in foveal vision and with small items, primarily involved VPL and the anterior pulvinar. The grasping deficits following pulvinar lesions are consistent with both human^23^ and monkey lesion studies^24,25^, and it is possible that subportions such as the anterior pulvinar portion are particularly important for this to occur.

### The central thalamus as the critical lesion site for optic ataxia

The critical lesion we found in optic ataxia patients was not restricted to one circumscribed region within the Morel atlas system. Instead, it encompassed several nuclei within and close to the internal medullary laminar (IML) complex. Interestingly, this region matches the so-called ‘central thalamus’, also sometimes called ‘oculomotor thalamus’. Since the pioneering electrophysiological studies in awake monkeys, it is considered to include the anterior group of the intralaminar nuclei (central lateral and central medial nucleus (CL, CM)), but also adjacent regions such as the medial portion of ventrolateral (VL) and the lateral portion of the mediodorsal thalamus (MD) ^39–41^. Thus, this ‘central thalamus’ region matches our OA-relevant thalamic lesion location quite well. In contrast to anatomical delineation schemes, the ‘central thalamus’ has been functionally defined by single neuron recordings in monkeys as a region where eye movement-related activity was observed ^42^. More generally, this ‘central thalamus’ region is considered a ‘higher-order’ nucleus complex that receives afferent inputs from the cerebellum and brainstem regions and connects to fronto-parietal regions involved in eye movement control ^41^. Neurons in this ‘central thalamus’ region fire in all visually guided task periods (cue, delay, saccade) and exhibit contralateral side preferences for visual saccade cues ^43^. Importantly, neurons in this central thalamus region also exhibit persistent activity as a function of eye position ^43,44^, as many sensorimotor cortical regions such as parietal cortex do ^45–48^. Strong support for the importance of the central thalamus to eye position perception is further provided by lesion studies in humans, showing impairments in perceiving and integrating eye position information across saccades ^49–51^. Eye position information is thought to be critical for transformations between retinal and egocentric limb/body coordinates that are relevant for visually guided reaches ^52^. Optic ataxia following lesions in this thalamic nucleus complex could then be construed as a failure to transmit correct eye position information to parietal cortex, which then impairs the integration of retinal, eye and hand signals as a prerequisite of reaching visual objects. Or not mutually exclusive, as a breakdown of parietal cortex capacity to integrate different directional eye- and hand-related information, thus depriving motor-related areas in prefrontal cortex of important visuomotor inputs^53^. Methods such as resting state MRI or lesion-based network mapping would be a good further step.

The lesion differences between OA and non-OA patients also revealed parts of the so called ‘motor thalamus’ such as the paralaminar portions of the central lateral (CL), ventral lateral (VL), mediodorsal (MD) and ventral posterior nucleus (VPL) ^54^. Consistent with previous studies, some patients who showed contralesional optic ataxia, also exhibited a mild decreased force in the MRC or hemiataxia in the finger-to-nose task in clinical testing. Previous electrophysiological studies have provided evidence from monkey studies that neurons in those nuclei increase their activity in visually guided reach tasks ^14,55^. How those neurons respond in tasks where gaze and movement targets are coordinated vs. dissociated, and whether those neurons contain information about eye position remains to be elucidated.

### Lack of optic ataxia following pulvinar lesions

To our great surprise, patients with lesions in the pulvinar did not typically exhibit optic ataxia. Based on its reciprocal connectivity to fronto-parietal regions that subserve visuomotor functions ^56–58^ and importantly to the optic ataxia-linked SPL/IPL regions ^8^, we had hypothesized that (dorsal) pulvinar lesions would preferentially lead to optic ataxia. Furthermore, there were two pulvinar inactivation/lesion studies in macaque and marmoset monkeys who reported ‘optic ataxia’ following pulvinar lesions ^24,30^. The first study in macaques inactivated the dorso-lateral portion of the pulvinar and reported imprecise and slowed reach-grasping along with failures to appropriately preshape the hand, most pronounced with the contralesional hand and space ^30^. Another study in marmosets showed that ablation of the ventral pulvinar in early life results in persistent misreaches and grasping deficits ^24^. Similar to the ‘central thalamus’ detailed above, neurons in ventral and dorsal pulvinar portions are strongly modulated by eye movements ^59,60^ and many pulvinar neurons carry information about (static) eye position during fixation and code visual stimuli in a dynamic spatial reference frame, from predominantly eye-centered visual encoding to final gaze representations in non-retinocentric (possibly body-centered) coordinates ^61^. The few studies which recorded pulvinar neurons in reach tasks in monkeys, reported neurons that either responded during reach planning, execution or post-reach ^62–64^. On the other hand, some basic reach properties of pulvinar neurons such as specificity for hand (left, right) and hand/space interactions and the underlying spatial reference frames remain unknown ^65^. It is not clear why damage in other higher-order thalamic nuclei (i.e. central thalamus) with similar connectivity should be more likely to induce optic ataxia. Since we excluded patients with visual field loss and thus LGN lesions, we have likely excluded patients with damage of the posterior choroidal artery that also supplies the more ventral portion of the pulvinar ^66^. Hence, our sample was biased towards more medial/anterior pulvinar portions. Further lesion studies should selectively also include more dorso-lateral pulvinar portions that have strong parietal connectivity as well ^67,68^.

### Limitations

Ischemic lesions are typically not restricted to a single nucleus but occur in combinations defined by the vascular supply ^66^. The anatomical specificity is thus somewhat limited by the stroke etiology in our patients, although the lesion subtraction method should largely correct for this ^69^. Unfortunately, the number of optic ataxia patients was too small to allow for statistically rigid voxel-based symptom-lesion mapping (VLBM) ^70,71^. Thus, future studies need to increase the number of thalamic stroke patients. Digital movement tracking with extraction of time courses would also have helped to quantify more subtle changes of reach-grasp behavior such as slowing. Since we did not systematically vary head and trunk position, we cannot say whether our patients had a specific problem with eye-based reference frame transformation or whether they had problems in other, i.e. head-based or trunk-based spatial reference frames ^46^. While we conducted computer-based visual tasks to control for general vision and spatial attention problems, we do not have a systematic assessment proprioceptive deficits. Future studies should also test whether the misreaches are specific for the visual modality or whether they would occur when pointing towards auditory or somatosensory targets ^72^. Since the critical thalamic lesion site for optic ataxia is distributed through several thalamic nuclei as defined by structural anatomy, a lesion-based network mapping (LSM) approach could be useful in future studies to understand the remote network effects better ^73–76^.

## Conclusion

We analysed 28 stroke patients with circumscribed thalamic lesions and detected five patients who resembled the well-known optic ataxia as observed in patients with parietal lesions. The critical lesion site appears to be a group of nuclei often referred to as ‘central thalamus’.

## Supporting information

Main Manuscript

## Data availability

Anonymized data may be shared on request to the corresponding author from a qualified investigator for non-commercial use, subject to restrictions according to participant consent and data protection legislation.

## Acknowledgements

We thank Severin Heumueller for excellent technical and computer support, Kristina Miloserdov and Maria Mendoza for participation in the patient testing. Sabine Nuhn we thank for detailed neuropsychological assessments. Britta Perl and Ilona Pfahlert we thank for assistance with the acquisition of the MRI data.

## Supplementary material

Supplementary material is available at *Brain* online.

## References

1. Blangero A, Gaveau V, Luaute J, et al. A hand and a field effect in on-line motor control in unilateral optic ataxia. Cortex. 2008;44(5):560–568. doi:10.1016/j.cortex.2007.09.004

2. Rossetti Y, Pisella L, McIntosh RD. Definition: Optic ataxia. Cortex. 2019;121:481. doi:10.1016/j.cortex.2019.09.004

3. Borchers S, Müller L, Synofzik M, Himmelbach M. Guidelines and quality measures for the diagnosis of optic ataxia. Front Hum Neurosci. 2013;7. doi:10.3389/fnhum.2013.00324

4. Perenin MT, Vighetto A. Optic ataxia: a specific disruption in visuomotor mechanisms. I. Different aspects of the deficit in reaching for objects. Brain. 1988;111 ( Pt 3):643–674.

5. Castiello U. The neuroscience of grasping. Nat Rev Neurosci. 2005;6(9):726–736. doi:10.1038/nrn1744

6. Andersen RA, Andersen KN, Hwang EJ, Hauschild M. Optic Ataxia: From Balint’s Syndrome to the Parietal Reach Region. Neuron. 2014;81(5):967–983. doi:10.1016/j.neuron.2014.02.025

7. Karnath HO, Perenin MT. Cortical Control of Visually Guided Reaching: Evidence from Patients with Optic Ataxia. Cerebral Cortex. 2005;15(10):1561–1569. doi:10.1093/cercor/bhi034

8. Gamberini M, Passarelli L, Filippini M, Fattori P, Galletti C. Vision for action: thalamic and cortical inputs to the macaque superior parietal lobule. Brain Struct Funct. 2021;226(9):2951–2966. doi:10.1007/s00429-021-02377-7

9. Yeterian EH, Pandya DN. Corticothalamic connections of the posterior parietal cortex in the rhesus monkey. J Comp Neurol. 1985;237(3):408–426. doi:10.1002/cne.902370309

10. Cappe C, Morel A, Rouiller EM. Thalamocortical and the dual pattern of corticothalamic projections of the posterior parietal cortex in macaque monkeys. Neuroscience. 2007;146(3):1371–1387. doi:10.1016/j.neuroscience.2007.02.033

11. Hwang K, Bertolero MA, Liu WB, D’Esposito M. The Human Thalamus Is an Integrative Hub for Functional Brain Networks. J Neurosci. 2017;37(23):5594–5607. doi:10.1523/JNEUROSCI.0067-17.2017

12. Kumar VJ, Beckmann CF, Scheffler K, Grodd W. Relay and higher-order thalamic nuclei show an intertwined functional association with cortical-networks. Commun Biol. 2022;5(1):1187. doi:10.1038/s42003-022-04126-w

13. Sommer MA. The role of the thalamus in motor control. Curr Opin Neurobiol. 2003;13(6):663–670. doi:10.1016/j.conb.2003.10.014

14. Bosch-Bouju C, Hyland BI, Parr-Brownlie LC. Motor thalamus integration of cortical, cerebellar and basal ganglia information: implications for normal and parkinsonian conditions. Front Comput Neurosci. 2013;7:163. doi:10.3389/fncom.2013.00163

15. Caplan LR, DeWitt LD, Pessin MS, Gorelick PB, Adelman LS. Lateral Thalamic Infarcts. Archives of Neurology. 1988;45(9):959–964. doi:10.1001/archneur.1988.00520330037008

16. Melo TP, Bogousslavsky J, Moulin T, Nader J, Regli F. Thalamic ataxia. J Neurol. 1992;239(6):331–337. doi:10.1007/BF00867590

17. Bastian AJ, Thach WT. Cerebellar outflow lesions: a comparison of movement deficits resulting from lesions at the levels of the cerebellum and thalamus. Ann Neurol. 1995;38(6):881–892. doi:10.1002/ana.410380608

18. Chen H, Hua SE, Smith MA, Lenz FA, Shadmehr R. Effects of Human Cerebellar Thalamus Disruption on Adaptive Control of Reaching. Cerebral Cortex. 2006;16(10):1462–1473. doi:10.1093/cercor/bhj087

19. Kagan I, Gibson L, Spanou E, Wilke M. Effective connectivity and spatial selectivity-dependent fMRI changes elicited by microstimulation of pulvinar and LIP. Neuroimage. 2021;240:118283. doi:10.1016/j.neuroimage.2021.118283

20. Halassa MM, Kastner S. Thalamic functions in distributed cognitive control. Nature Neuroscience. 2017;20(12):1669–1679. doi:10.1038/s41593-017-0020-1

21. Fiebelkorn IC, Pinsk MA, Kastner S. The mediodorsal pulvinar coordinates the macaque fronto-parietal network during rhythmic spatial attention. Nature Communications. 2019;10(1):215. doi:10.1038/s41467-018-08151-4

22. Frassinetti F, Bonifazi S, Làdavas E. The influence of spatial coordinates in a case of an optic ataxia-like syndrome following cerebellar and thalamic lesion. Cognitive Neuropsychology. 2007;24(3):324–337. doi:10.1080/02643290701275857

23. Wilke M, Schneider L, Dominguez-Vargas AU, et al. Reach and grasp deficits following damage to the dorsal pulvinar. Cortex. 2018;99:135–149. doi:10.1016/j.cortex.2017.10.011

24. Mundinano IC, Fox DM, Kwan WC, et al. Transient visual pathway critical for normal development of primate grasping behavior. Proc Natl Acad Sci USA. 2018;115(6):1364–1369. doi:10.1073/pnas.1717016115

25. Wilke M, Turchi J, Smith K, Mishkin M, Leopold DA. Pulvinar Inactivation Disrupts Selection of Movement Plans. Journal of Neuroscience. 2010;30(25):8650–8659. doi:10.1523/JNEUROSCI.0953-10.2010

26. Fels M, Geissner E. Neglect-Test (NET): ein Verfahren zur Erfassung visueller Neglectphänomene; Handanweisung; deutsche überarbeitete Adaption des Behavioural Inattention Test (Wilson, Cockburn & Halligan, 1987). Published online 1997.

27. Bickerton WL, Samson D, Williamson J, Humphreys GW. Separating forms of neglect using the Apples Test: validation and functional prediction in chronic and acute stroke. Neuropsychology. 2011;25(5):567–580. doi:10.1037/a0023501

28. Posner MI, Snyder CR, Davidson BJ. Attention and the detection of signals. J Exp Psychol. 1980;109(2):160–174.

29. Rengachary J, He BJ, Shulman GL, Corbetta M. A behavioral analysis of spatial neglect and its recovery after stroke. Front Hum Neurosci. 2011;5:29. doi:10.3389/fnhum.2011.00029

30. Wilke M, Turchi J, Smith K, Mishkin M, Leopold DA. Pulvinar Inactivation Disrupts Selection of Movement Plans. Journal of Neuroscience. 2010;30(25):8650–8659. doi:10.1523/JNEUROSCI.0953-10.2010

31. Morel A, Magnin M, Jeanmonod D. Multiarchitectonic and stereotactic atlas of the human thalamus. J Comp Neurol. 1997;387(4):588–630. doi:10.1002/(sici)1096-9861(19971103)387:4<588::aid-cne8>3.0.co;2-z

32. Krauth A, Blanc R, Poveda A, Jeanmonod D, Morel A, Székely G. A mean three-dimensional atlas of the human thalamus: generation from multiple histological data. Neuroimage. 2010;49(3):2053–2062. doi:10.1016/j.neuroimage.2009.10.042

33. Crawford JR, Garthwaite PH. Investigation of the single case in neuropsychology: confidence limits on the abnormality of test scores and test score differences. Neuropsychologia. 2002;40(8):1196–1208. doi:10.1016/s0028-3932(01)00224-x

34. Rossetti Y, Pisella L, Vighetto A. Optic ataxia revisited: visually guided action versus immediate visuomotor control. Experimental brain research Experimentelle Hirnforschung Experimentation cerebrale. 2003;153(2):171–179. doi:10.1007/s00221-003-1590-6

35. Carey DP, Coleman RJ, Sala SD. Magnetic Misreaching. Cortex. 1997;33(4):639–652. doi:10.1016/S0010-9452(08)70722-6

36. Jackson SR, Newport R, Mort D, Husain M. Where the eye looks, the hand follows; limb-dependent magnetic misreaching in optic ataxia. Curr Biol. 2005;15(1):42–46. doi:10.1016/j.cub.2004.12.063

37. Vindras P, Blangero A, Ota H, Reilly KT, Rossetti Y, Pisella L. The Pointing Errors in Optic Ataxia Reveal the Role of “Peripheral Magnification” of the PPC. Front Integr Neurosci. 2016;10. doi:10.3389/fnint.2016.00027

38. Cavina-Pratesi C, Ietswaart M, Humphreys GW, Lestou V, Milner AD. Impaired grasping in a patient with optic ataxia: Primary visuomotor deficit or secondary consequence of misreaching? Neuropsychologia. 2010;48(1):226–234. doi:10.1016/j.neuropsychologia.2009.09.008

39. Schlag J, Schlag-Rey M, Peck CK, Joseph JP. Visual responses of thalamic neurons depending on the direction of gaze and the position of targets in space. Exp Brain Res. 1980;40(2):l70–84.

40. Schlag J, Schlag-Rey M. Role of the central thalamus in gaze control. Prog Brain Res. 1986;64:191–201. doi:10.1016/S0079-6123(08)63413-5

41. Tanaka M, Kunimatsu J. Contribution of the central thalamus to the generation of volitional saccades. Eur J Neurosci. 2011;33(11):2046–2057. doi:10.1111/j.1460-9568.2011.07699.x

42. Schlag-Rey M, Schlag J. Visuomotor functions of central thalamus in monkey. I. Unit activity related to spontaneous eye movements. J Neurophysiol. 1984;51(6):1149–1174. doi:10.1152/jn.1984.51.6.1149

43. Wyder MT, Massoglia DP, Stanford TR. Quantitative Assessment of the Timing and Tuning of Visual-Related, Saccade-Related, and Delay Period Activity in Primate Central Thalamus. Journal of Neurophysiology. 2003;90(3):2029–2052. doi:10.1152/jn.00064.2003

44. Tanaka M. Spatiotemporal Properties of Eye Position Signals in the Primate Central Thalamus. Cerebral Cortex. 2007;17(7):1504–1515. doi:10.1093/cercor/bhl061

45. DeSouza JF, Dukelow SP, Gati JS, Menon RS, Andersen RA, Vilis T. Eye position signal modulates a human parietal pointing region during memory-guided movements. J Neurosci. 2000;20(15):5835–5840. doi:10.1523/JNEUROSCI.20-15-05835.2000

46. Cohen YE, Andersen RA. A common reference frame for movement plans in the posterior parietal cortex. Nature reviews Neuroscience. 2002;3(7):553–562. doi:10.1038/nrn873

47. Wang X, Zhang M, Cohen IS, Goldberg ME. The proprioceptive representation of eye position in monkey primary somatosensory cortex. Nat Neurosci. 2007;10(5):640–646. doi:10.1038/nn1878

48. McFadyen JR, Heider B, Karkhanis AN, et al. Robust Coding of Eye Position in Posterior Parietal Cortex despite Context-Dependent Tuning. J Neurosci. 2022;42(20):4116–4130. doi:10.1523/JNEUROSCI.0674-21.2022

49. Gaymard B, Rivaud S, Pierrot-Deseilligny C. Impairment of extraretinal eye position signals after central thalamic lesions in humans. Exp Brain Res. 1994;102(1):1–9.

50. Ostendorf F, Liebermann D, Ploner CJ. Human thalamus contributes to perceptual stability across eye movements. Proceedings of the National Academy of Sciences. 2010;107(3):1229–1234. doi:10.1073/pnas.0910742107

51. Ostendorf F, Liebermann D, Ploner CJ. A role of the human thalamus in predicting the perceptual consequences of eye movements. Front Syst Neurosci. 2013;7:10. doi:10.3389/fnsys.2013.00010

52. Andersen RA, Snyder LH, Li CS, Stricanne B. Coordinate transformations in the representation of spatial information. Curr Opin Neurobiol. 1993;3(2):171–176.

53. Battaglia-Mayer A, Caminiti R. Optic ataxia as a result of the breakdown of the global tuning fields of parietal neurones. Brain. 2002;125(Pt 2):225–237.

54. Prevosto V, Sommer MA. Cognitive control of movement via the cerebellar-recipient thalamus. Front Syst Neurosci. 2013;7:56. doi:10.3389/fnsys.2013.00056

55. van Donkelaar P, Stein JF, Passingham RE, Miall RC. Temporary inactivation in the primate motor thalamus during visually triggered and internally generated limb movements. J Neurophysiol. 2000;83(5):2780–2790. doi:10.1152/jn.2000.83.5.2780

56. Gutierrez C, Cola MG, Seltzer B, Cusick C. Neurochemical and connectional organization of the dorsal pulvinar complex in monkeys. Journal of Comparative Neurology. 2000;419(1):61–86.

57. Kaas JH, Lyon DC. Pulvinar contributions to the dorsal and ventral streams of visual processing in primates. Brain Res Rev. 2007;55(2):285–296.

58. Arcaro MJ, Pinsk MA, Chen J, Kastner S. Organizing principles of pulvino-cortical functional coupling in humans. Nature Communications. 2018;9(1):5382. doi:10.1038/s41467-018-07725-6

59. Benevento LA, Port JD. Single neurons with both form/color differential responses and saccade-related responses in the nonretinotopic pulvinar of the behaving macaque monkey. Vis Neurosci. 1995;12(3):523–544. doi:10.1017/s0952523800008439

60. Schneider L, Dominguez-Vargas AU, Gibson L, Wilke M, Kagan I. Visual, delay, and oculomotor timing and tuning in macaque dorsal pulvinar during instructed and free choice memory saccades. Cereb Cortex. 2023;33(21):10877–10900. doi:10.1093/cercor/bhad333

61. Schneider L, Dominguez-Vargas AU, Gibson L, Kagan I, Wilke M. Eye position signals in the dorsal pulvinar during fixation and goal-directed saccades. J Neurophysiol. 2020;123(1):367–391. doi:10.1152/jn.00432.2019

62. Cudeiro J, González F, Pérez R, Alonso JM, Acuña C. Does the pulvinar-LP complex contribute to motor programming? Brain Res. 1989;484(1-2):367–370. doi:10.1016/0006-8993(89)90383-1

63. Acuña C, Cudeiro J, Gonzalez F, Alonso JM, Perez R. Lateral-posterior and pulvinar reaching cells--comparison with parietal area 5a: a study in behaving Macaca nemestrina monkeys. Exp Brain Res. 1990;82(1):158–166. doi:10.1007/BF00230847

64. Grieve KL, Acuña C, Cudeiro J. The primate pulvinar nuclei: vision and action. Trends Neurosci. 2000;23(1):35–39. doi:10.1016/s0166-2236(99)01482-4

65. Wilke M, Kagan I. Visuospatial and motor deficits following pulvinar lesions. In: The Cerebral Cortex and Thalamus. Martin Usrey and S. Murray Sherman. Oxford University Press; 2024:764–775.

66. Schmahmann JD. Vascular Syndromes of the Thalamus. Stroke. 2003;34(9):2264–2278. doi:10.1161/01.STR.0000087786.38997.9E

67. Benarroch EE. Pulvinar Associative role in cortical function and clinical correlations. Neurology. 2015;84(7):738–747.

68. Froesel M, Cappe C, Ben Hamed S. A multisensory perspective onto primate pulvinar functions. Neurosci Biobehav Rev. 2021;125:231–243. doi:10.1016/j.neubiorev.2021.02.043

69. Rorden C, Karnath HO. Using human brain lesions to infer function: a relic from a past era in the fMRI age? Nat Rev Neurosci. 2004;5(10):813–819. doi:10.1038/nrn1521

70. Bates E, Wilson SM, Saygin AP, et al. Voxel-based lesion-symptom mapping. Nat Neurosci. 2003;6(5):448–450. doi:10.1038/nn1050

71. Sperber C, Karnath HO. On the validity of lesion-behaviour mapping methods. Neuropsychologia. 2018;115:17–24. doi:10.1016/j.neuropsychologia.2017.07.035

72. Jackson SR, Newport R, Husain M, Fowlie JE, O’Donoghue M, Bajaj N. There may be more to reaching than meets the eye: Re-thinking optic ataxia. Neuropsychologia. 2009;47(6):1397–1408. doi:10.1016/j.neuropsychologia.2009.01.035

73. Karnath HO, Ferber S, Dichgans J. The neural representation of postural control in humans. Proc Natl Acad Sci USA. 2000;97(25):13931–13936. doi:10.1073/pnas.240279997

74. Karnath HO, Sperber C, Rorden C. Mapping human brain lesions and their functional consequences. Neuroimage. 2018;165:180–189. doi:10.1016/j.neuroimage.2017.10.028

75. Rosenzopf H, Klingbeil J, Wawrzyniak M, et al. Thalamocortical Networks Involved in Pusher Syndrome. Neuroscience; 2022. doi:10.1101/2022.10.12.511887

76. Stockert A, Hormig-Rauber S, Wawrzyniak M, et al. Involvement of Thalamocortical Networks in Patients With Poststroke Thalamic Aphasia. Neurology. 2023;100(5). doi:10.1212/WNL.0000000000201488

