## Supplementary material for "Optic Ataxia in Patients with Thalamic Lesions": Main Manuscript

#these authors contributed equally

**Supplementary material includes:**

Supplementary Methods

Single Patient Descriptions

7 Supplementary Figures

4 Supplementary Tables

### Supplementary Methods

#### Testing for spatial neglect

Testing for spatial neglect was conducted with several paper and pencil tests of the German neglect battery [1]: (i) line bisection (ii) line cancellation (iii) star cancellation and the Apples test [2]. All these paper-based tests were presented on a horizontally-oriented DIN A4 sheet. Time of test completion depended on the patient indicating to be satisfied with his/her performance. In order to assess the spatial bias we calculated the center of cancellation (CoC) with custom-written software similarly to [3]. In short, the CoC is a continuous measure that represents the mean horizontal coordinate of the detected items, first in terms of centimeters and then transformed into a normalized score [4], based on the performance of our group of healthy controls. Apart from the paper and pencil tasks, visual attention deficits were tested with the Posner paradigm, shown to be sensitive even for patients with mild neglect [5,6]. The task design and temporal structure of the Posner task was adapted from previous neglect studies [7]. In the computerized paradigm which we adopted from [7], subjects detect a visual stimulus presented to the left or right from fixation. The side of attention is cued by an additional stimulus that appears before the target appearance. Critically, in the majority of trials (typically 75%, 'valid'), the spatial cue correctly predicts the side of target appearance, while pointing towards the opposite hemifield in the remaining trials (typically 25%, 'invalid'). Three key parameters were derived from the Posner-Tak: i) Longer detection latencies (and/or a higher number of misses) for contralesional items indicate deficits in visual perception/attention ('side/visual field effect'); ii) Larger response delays between trials with valid cues vs. invalidly cued location indicate deficits in re-orienting attention ('validity effect'); iii) Unproportionally longer response delays for contralesional targets when the ipsilesional side was cued (i.e. most likely attended) are taken as deficits to disengage attention ('disengagement effect'). For our neglect assessment we used the reaction times for contralesional targets. A z-score smaller than -2 in more than 2 of the 5 tasks (paper and pencil tests and Posner task) was considered as spatial neglect.

#### Assessment of grasping performance

We assessed fine visually-guided reaching-grasping abilities of the patients and the control group by letting them pick small objects ('Smarties' candies, oblate spheroids with a minor axis of about 5 mm and a major axis of about 12 mm) from the center and peripheral table positions. The task and scoring was adopted to be comparable to our previous pulvinar lesion studies in non-human primates and humans[8,9] (**Supplementary Fig. 1**). Participants were always allowed to look at the objects. We calculated an error score (fluent (0), slowed (1), corrected during (2) / after the first (3) or the second (4) attempt, not successful (5). Subjects performed 36 (median) reach-grasp trials per session with either the left or right hand. Healthy subjects rarely made any reach-grasp errors while grasping the objects independently of hand or spatial position, yielding a median error score of 0. As a group, patients with thalamic lesions exhibited impaired reach-grasp movements, more often for the contralesional hand and space (Mann-Whitney U test, $U=273.5$, $p<0.001$), but significant also for the ipsilesional hand and space ($U=410$, $p=0.049$).

#### Anatomical lesion mapping

Anatomical data were analysed using the FMRIB software library (FSL 5.0.7, Center for Functional Magnetic Resonance Imaging of the Brain, University of Oxford, UK www.fmrib.ox.ac.uk/fsl). Procedures have been described before [23]. The T1-weighted tFLASH dataset was skull stripped (brain extraction tool BET) and registered to the standard brain template of the Montreal Neurological Institute at a 1 mm isotropic resolution (MNI152, provided with FSL), using the FMRIB's linear registration tool (FLIRT: 12 parameter affine transformation). The resulting transformation matrix was applied to the whole-head tFLASH dataset (without skull stripping) and the T2-weighted FLAIR dataset as well, after the FLAIR images had been coregistered to the T1 images of the same subject. In a second step, the linearly coregistered, whole-head tFLASH dataset was non-linearly registered to the MNI152 template using the FMRIB's non-linear registration tool (FNIRT). Again, the resulting transformation values were applied to the FLAIR dataset. Finally, FLIRT was used to sample the data to 0.5 mm isotropic resolution using the MNI152 template at 0.5 mm resolution. A digitized version of the Morel atlas of the thalamus [31] provided by Krauth et al. [32], registered to the high-resolution MNI 152 template allowed the visualization of the thalamic nuclei in respect to the thalamic lesions, which were manually segmented on FLAIR MR images using MRIcron and transformed to MNI space (via coregistration of individual FLAIR images to individual T1-weighted images followed by normalization to the MNI152 T1 0.5mm template using FSL FLIRT, FNIRT). Individual scans and results of the lesion mapping are shown in **Supplementary Fig. 2 and Supplementary Table 1.**

### Single Patient Descriptions

#### Patient N054: Optic Ataxia without Grasping deficit

N054 is a 70-year old, right handed man who suffered from a left thalamic ischemic stroke. The patient presented with difficulties to find words. The **MRI** showed a lesion in the left medial (MD, CeM, CL, CM) and lateral thalamus (VA, VL, VM, VP). **Neurological examination** at admission showed a modest non-fluent aphasia. The patient's cranial nerve examination showed a facial asymmetry to the right. The reflex-status, muscle tone and strength were normal. **There was no sign for ataxia with a metric finger-to-nose test on both sides.** The experimental tasks and MRI's were performed within the first week after hospital admission due to symptom onset. **Neurophysiological** examinations showed an afferent deficit from both arms and legs as well as an efferent deficit to both legs. **Neuropsychological** assessment revealed normal executive, but abnormal memory and language functions. The Mini Mental Status was lower than average with 24/30 points. Oculomotor functions were normal and visual fields were intact. There was no sign of spatial neglect or extinction. EEG was normal with an alpha ground rhythm. **Grasping:** N054 showed normal reach-grasp scores in the smarties test with both hands (Grasping CH-CS: z = 0.43, IH-IS: z = 0.80). **Optic Ataxia:** In the optic ataxia task he mostly made corrected or uncorrected errors in the non-foveal condition with the right (i.e. contralesional) hand (corrected: 5.3%, uncorrected: 13.2%). However, the majority of trials (81.6%) with this hand were rated effective and fluent. The slowed and corrected reaches with the right hand in the right space led to a pathological optic ataxia score for the contralesional hand/space in this patient (z = -3.67). The OA scores for the IH-IS condition were in the lower normal range (z = -1.57). While most errors occurred in the combination contralesional hand/field (CH-CS) condition, no other clear hand or space effect can be observed.

#### Patient N056: Optic Ataxia without Grasping deficit

N056 is a 24-year old, right handed man who suffered from a bilateral (left > right) thalamic hemorrhagic stroke resulting from a sinus venous thrombosis. The patient complained about headaches and fatigue at admission. The **MRI** showed a lesion in the left medial (MD, CL, CM) and lateral thalamus (VL, VP). On the right side, there was a smaller VA damage. **Neurological examination** at admission showed a general slowness as well as a slight paresis and fine motor difficulties of the right arm. Sensibility to touch was intact on both sides. The patient's cranial nerve examination was normal. The reflex-status, muscle tone and other sensory tests were normal. Oculomotor functions were normal and visual fields were intact. There was no sign of spatial neglect or extinction. **There was no sign for ataxia with a metric finger-to-nose test on both sides.** **Neuropsychological** assessment revealed normal executive, normal memory and normal language functions. The Mini Mental Status was also normal with 30/30 points. The experimental tasks and MRI's were performed within two weeks after hospital admission due to symptom onset. EEG before testing was normal with an alpha ground rhythm. **Grasping:** N056 showed normal reach-grasp scores in the smarties test with both hands (Grasping CH-CS: $z = 0.43$, IH-IS: $z = 0.07$). **Optic Ataxia:** In the optic ataxia task he mostly made corrected or uncorrected errors in the non-foveal condition with the right hand (corrected: 16.6%, uncorrected: 11.1%). However, the majority of trials (72.2%) with this hand were rated effective and fluent. The slowed and corrected reaches with the right hand in the right space led to a pathological optic ataxia score for the contralesional hand/space in this patient ($z = -3.68$). The OA scores for the IH-IS condition were normal. Gross (i.e. uncorrected) reach errors occurred all away from the body and deviated in 4 out of 5 cases towards the right (i.e. contralesional) side. Most errors occurred in the combination contralesional hand/contralesional field (CH-CS) condition and he made also relatively more errors in the ipsilesional hand/contralesional field (IH-CS). There thus seems to be some space effect.

#### Patient N063: Optic Ataxia + Grasping deficit

N063 is a 50-year old, right handed woman who suffered from a bilateral thalamic ischemic stroke. The patient presented with dysaesthesia, i.e. tingling of the right arm and face. The **MRI** showed a bilateral lesion (left > right) encompassing the medial (CM, CL) and lateral thalamus (bilateral VP, but VL and VM on the left side only), and posterior thalamus (anterior and medial Pulvinar and LP). While the right thalamic lesion was visible in the MRI, it remained clinically silent. Thus for consistency, the left side is used as reference for the label 'contra- or ipsilesional' (i.e. right is contralesional). **Neurological examination** at admission showed a latent paresis on the right side (sinking of the arm). **The finger-to-nose test was slightly dysmetric on the right side.** The patient's cranial nerve examination was normal. The reflex-status, muscle tone and other sensory and muscle strength tests were normal. Oculomotor functions were normal and visual fields were intact. The experimental tasks and MRI's were performed within two weeks after hospital admission due to symptom onset. EEG was normal. **Neuropsychological** exam was conducted on day 8 after admission and revealed normal executive, normal memory and normal language functions. The Mini Mental Status was also normal with 30/30 points. There was no indication of spatial neglect or extinction. **Grasping:** N063 showed pathological reach-grasp scores in the smarties test with the contralesional (right) hand only (z = -6.9). Z-scores for grasping with the ipsilesional hand were normal (z = 0.21). **Optic Ataxia:** In the optic ataxia task she made a large amount of corrected errors in the non-foveal condition with the right hand. Only 30.9% trials with this hand were rated effective and fluent, while the remaining trials were slowed and ineffective but were corrected in flight. The slowed and corrected reaches with the right hand in the right space led to a pathological optic ataxia score in this patient (z = -4.6). The ipsi hand/field OA condition was not impaired (OA z-score: -1.32). Reach errors mainly occurred all away from the body and deviated towards the right (i.e. contralesional) side. While most errors occurred in the combination contralesional hand/field (CH-CS) condition, no other clear hand or space effect can be observed.

#### Patient N075 with small grasping deficits

N075 is a 63-year old, right handed man who suffered from a right thalamic ischemic stroke. The patient presented with hypaesthesia of the left cheek and hand. Neurological examination at admission confirmed a brachiofacial hypaesthesia on the left and found a facial asymmetry. The **MRI s**howed a primarily right sided lesion in the right lateral thalamus (VP, VL) with involvement of the medial group (CL, CM, MD (< 10%)) and anterior pulvinar. In the left thalamus, smaller (possibly older?) lesions in the medial (MD, CM, CL), VP and anterior pulvinar. **Neurological examination** showed that apart from the hypaesthesia, the patient's cranial nerve examination was elsewise normal. The reflex-status, muscle tone and other sensory tests were normal. Oculomotor functions were normal and visual fields were intact. There was no sign of spatial neglect or extinction, although he showed a lateralized deficit in the line bisection task. **There was no sign for ataxia with a metric finger-to-nose test on both sides. Neuropsychological assessments** were not performed. The **EEG** showed a normal alpha ground rhythm with intermittent fronto-temporal delta activity on the left side. **Grasping:** N075 showed borderline abnormal reach-grasp scores in the smarties test with the contralesional (left) hand (Grasping CH-CS: z = -1.81, IH-IS: z = -1.1). **Optic Ataxia:** In the optic ataxia task he mostly made corrected or uncorrected errors in the non-foveal condition with the contralesional hand (corrected: 3.6%, uncorrected: 14.29%). However, the majority of trials (82.1%) with this hand were rated effective and fluent. He had a pathological optic ataxia score for the contralesional (left) hand/space (z = -2.81). The OA scores for the IH-IS condition were normal (z = 0.71). Most errors occurred in the combination contralesional hand/contralesional field (CH-CS) condition and he also made relatively more errors in the ipsilesional hand/contralesional field (IH-CS). There thus seems to be some (contralesional) space effect.

#### Patient N093 with grasping deficits

N093 is a 71-year old, right handed man who suffered from a right thalamic ischemic stroke. The **MRI** showed a thalamic lesion that included the right medial thalamus (MD, CeM, CMm CL) as well as the lateral thalamus (VL, VM, VP). **Neurological examination** showed a brachiofacial paresis of the left side with a pronation of the left arm and dysarthria. The patient's cranial nerve examination was elsewise normal. The reflex-status, muscle tone and other sensory and muscle strength tests were normal. Sensibility to touch was intact on both sides. Oculomotor functions were normal and visual fields were intact. There was no sign of spatial neglect or extinction. **There was no sign for ataxia with a metric finger-to-nose test on both sides.** The Mini Mental Status was also normal with 30/30 points. **EEG** was normal with an alpha ground rhythm. **Grasping:** N093 showed impaired grasping in the smarties test with both hands (Grasping CH-CS: z = -3.82, IH-IS: z = 3.39). **Optic Ataxia:** In the optic ataxia task he mostly made corrected errors with his left (contralesional) hand (CH-CS: corrected: 23.3%, uncorrected: 3.3%). There was a clear hand effect, as he showed pathological error scores with the contralesional hand also when the movement was made into the ipsilesional space (CH-CS: corrected: 25.0%, uncorrected: 4.2%). Gross reach errors were mostly away from the body (5 away/ 2 towards) and deviated mostly towards the left side.

### Supplementary Figures

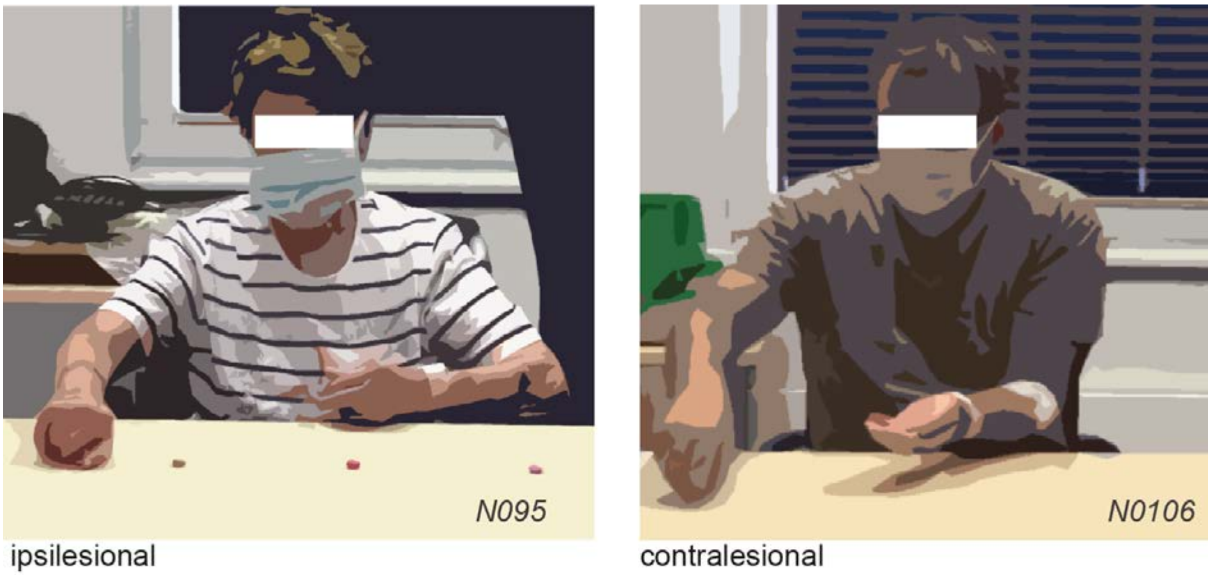


**Supplementary Figure 1 Reach-grasp task with small objects (smarties) in two example patients (drawing from video snapshot) with pathological grasp error scores (N095 and N106).** Typical grasp postures. Left panel: note the abnormal wrist angle and the absence of a precision grip. Right panel: note the grasp deviation from the object location.

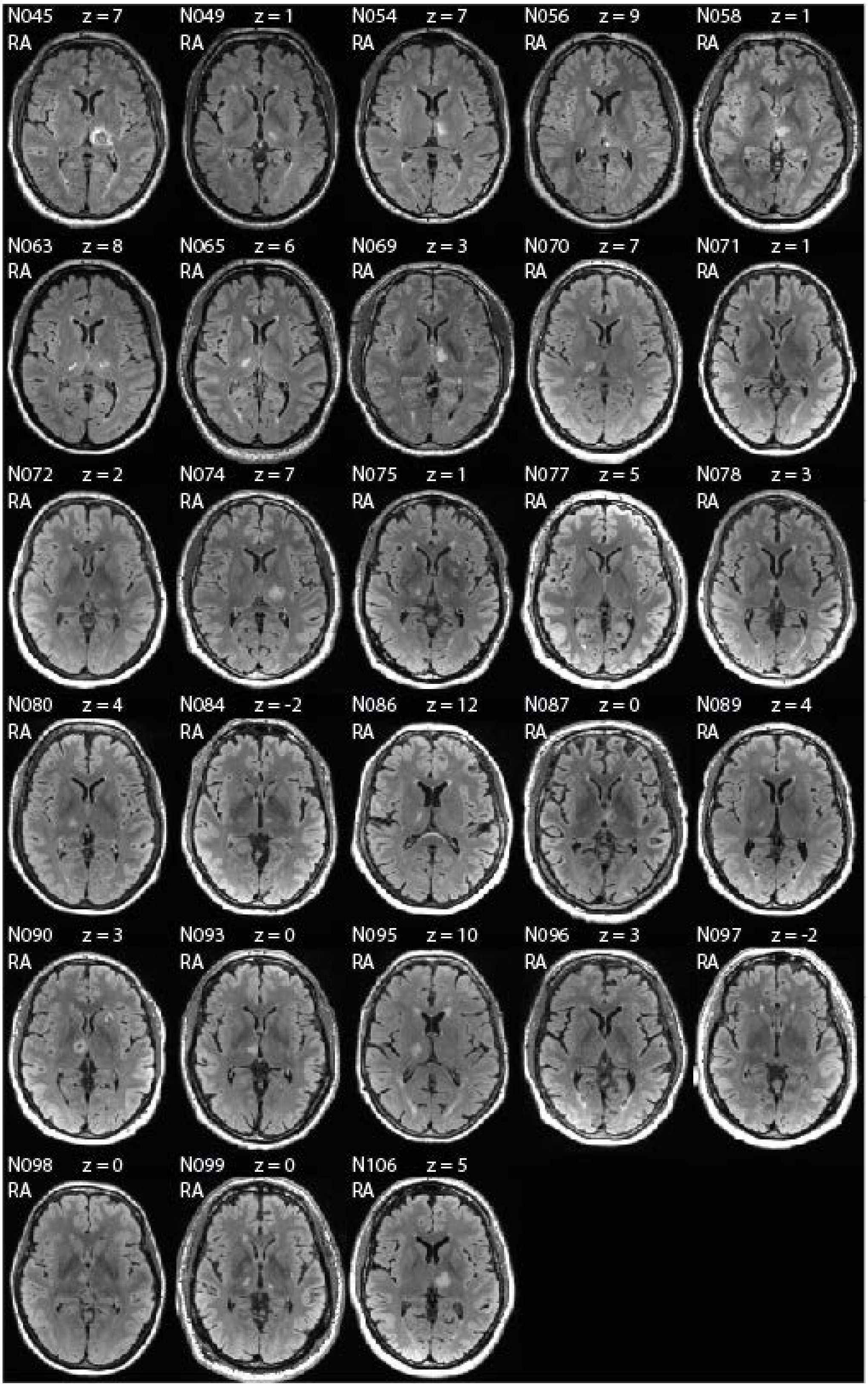


**Supplementary Figure 2 Lesion locations.** Reconstructed FLAIR axial images showing the lesion location for each patient. Lesion-centered axial slices are in MNI space, z-coordinates and patient abbreviation are reported on top of each slide. Slices are oriented in radiological convention: RA = right anterior, on the left.

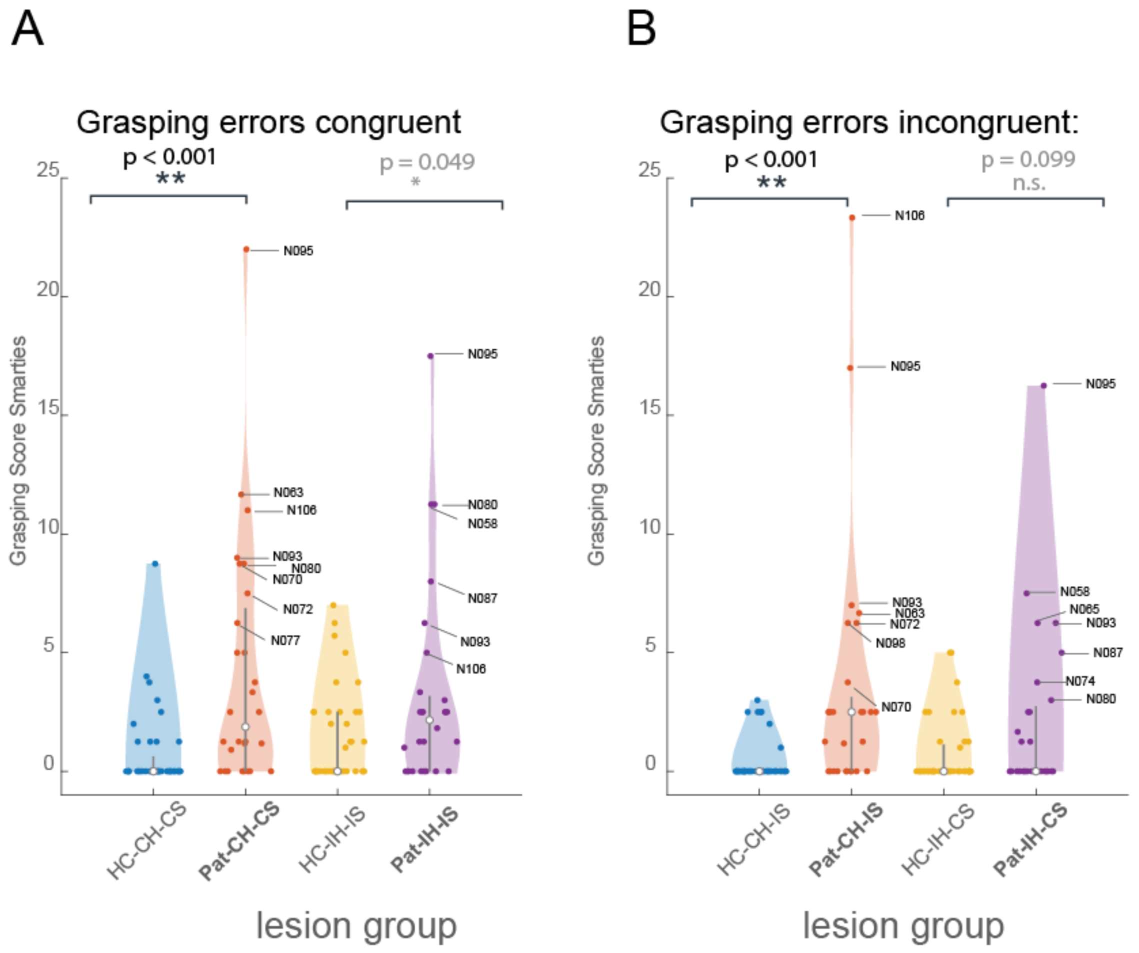


**Supplementary Figure 3 Grasping performance of the healthy controls and patients as a function of hand and space ($n_{patients}$ = 28; $n_{healthy}$ = 40).** Subjects were always allowed to look at the smarties before and during the reach-grasp. **(A)** Grasping errors across different hand-space thalamic patients and healthy controls. Congruent condition refers to the same hand and space (contralesional hand and space: CH-CS and ipsilesional hand and space: IH-IS). **(B)** Grasping error scores for the incongruent condition refer to contralesional hand and ipsilesional space: CH-IS or ipsilesional hand and contralesional space: IH-CS. Number of errors corresponds to the percentage of trials with corrected or uncorrected reach errors. The higher the number, the higher the percentage of trials with an error. Each dot represents the error score of each subject. The white dot in each violin plot represents the median of the specific subject group and the gray lines are the confidence intervals.

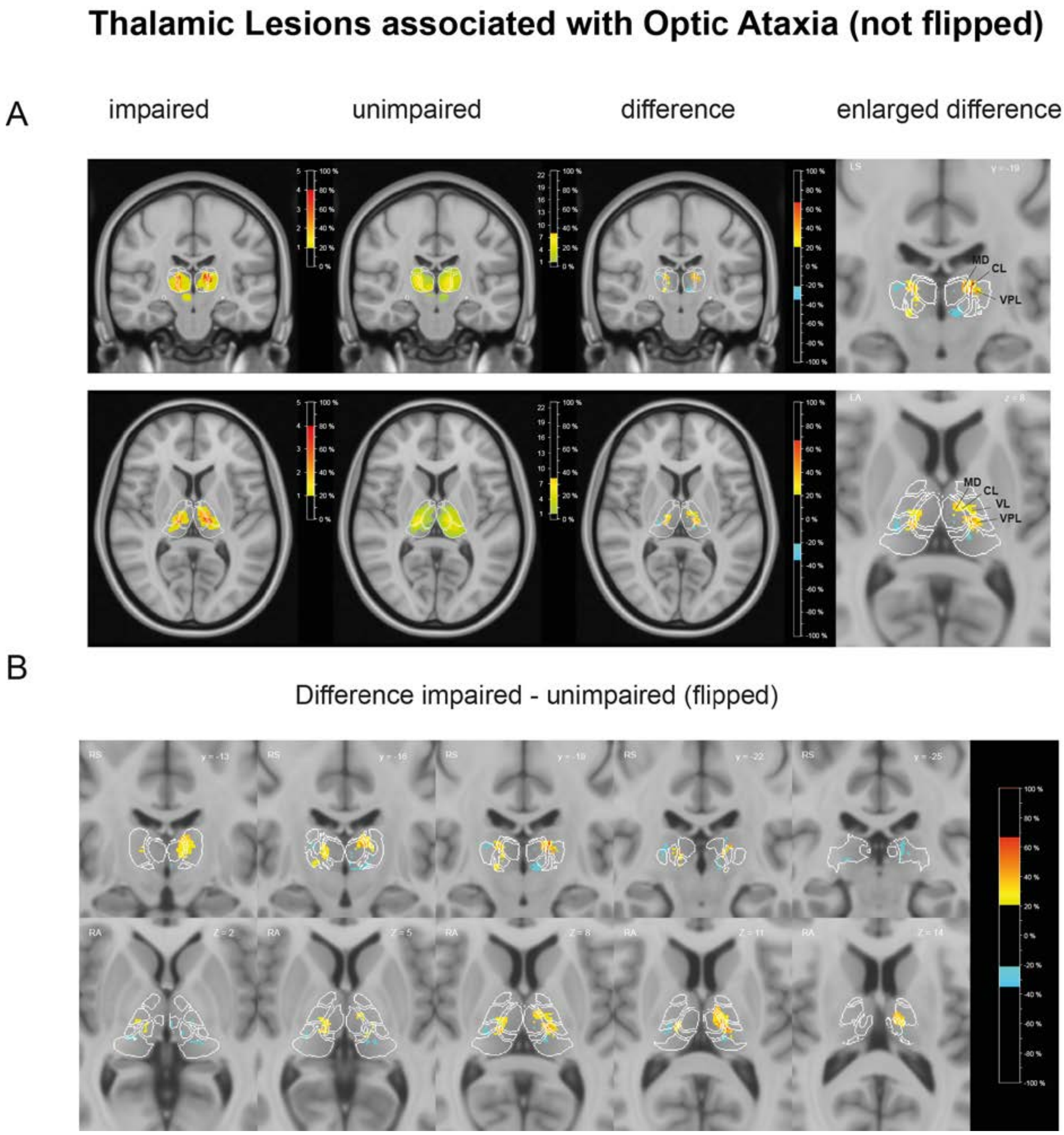


**Supplementary Figure 4 Lesion subtraction plots of the patients with and without optic ataxia. (A)** Lesion overlap of the patients with and without optic ataxia. **(B)** Percentage of overlapping lesions of the optic ataxia patients after subtraction of the patients without optic ataxia. The colorbar specifies the percentage of patients with lesions in each voxel, with blue indicating the fewest, and the hot colors (orange-red) a higher number of patients with an OA deficit. White outlines represent the borders of the lateral, medial, and posterior groups of thalamic nuclei as defined in the digital version of the Morel-atlas normalized to the 0.5 mm MNI152 T1 template. Colored schemes of nuclei outlines are depicted in **Supplementary Figure 5.**

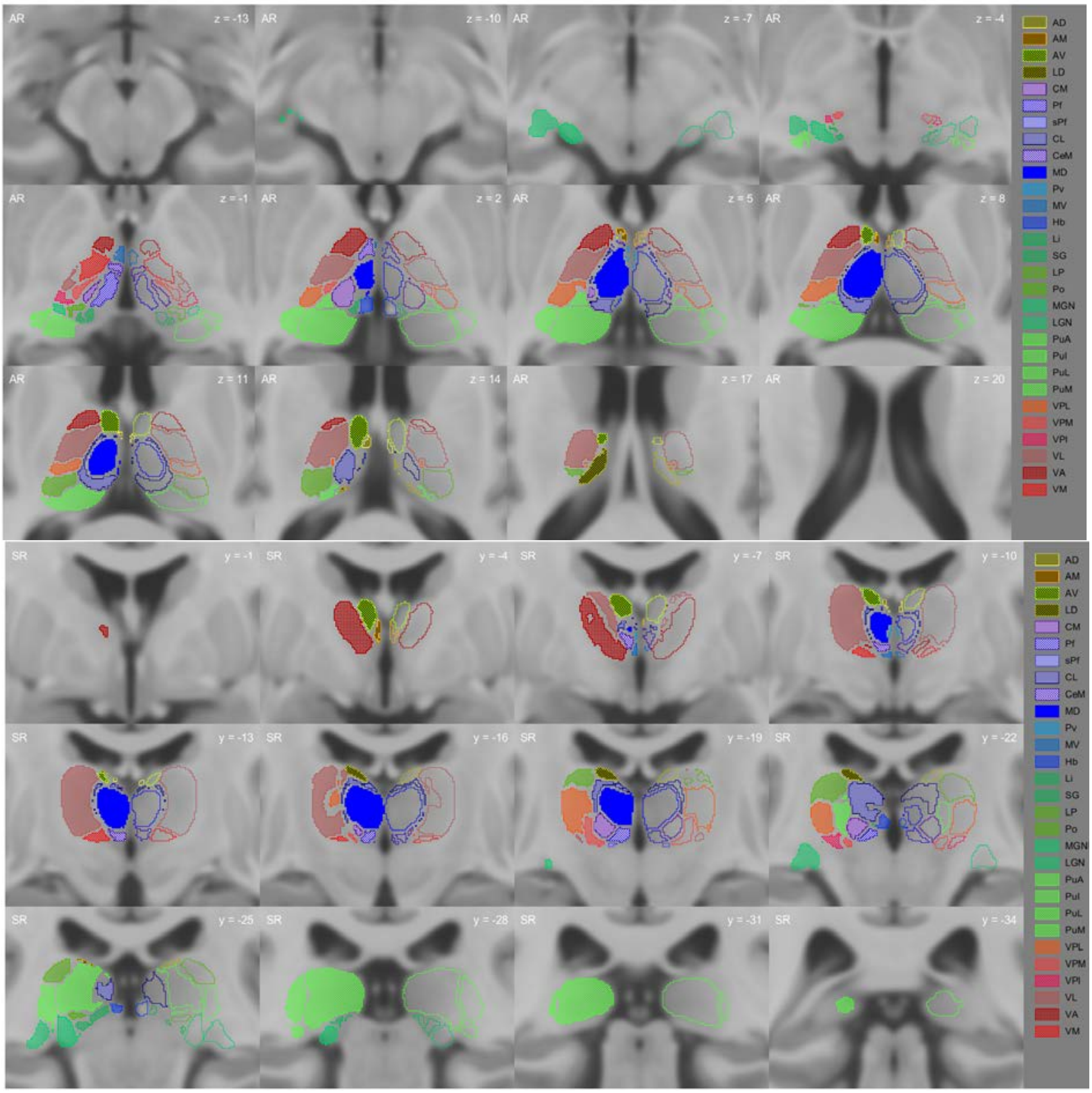


**Supplementary Figure 5 Coronal and Axial Slices with thalamic lesion labeling.** Abbreviations: **AD**, Anterior dorsal nucleus; **AM**, Anterior medial nucleus; **AV**, Anterior ventral nucleus; **LD**, Lateral dorsal nucleus; **MD**, Mediodorsal nucleus, **CM**, Centromedian nucleus; **Pf**, Parafascicular nucleus; **sPf**, Subparafascicular nucleus; **CL**, Central lateral nucleus; **CeM**, Central medial nucleus; **Pv**, Paraventricular nucleus; **MV**, Medioventral nucleus; **Hb**, Habenular nucleus; **PuA**, Anterior pulvinar; **PuI**, Inferior pulvinar; **PuL**, Lateral pulvinar; **PuM**, Medial pulvinar; **LP**, Lateral posterior nucleus; **Li**, Limitans nucleus; **Po**, Posterior nucleus; **SG**, Suprageniculate nucleus; **LGN**, Lateral geniculate nucleus; **MGN**, Medial geniculate nucleus; **VA**, Ventral anterior nucleus; **VL**, Ventral lateral nucleus; **VM**, Ventral medial nucleus; **VP**, Ventral posterior nucleus (**VPL/VPM/VPI**: lateral/medial/inferior). Thalamic nuclei impacted by the lesion were classified according to the Morel's neuroanatomical atlas [10,11].

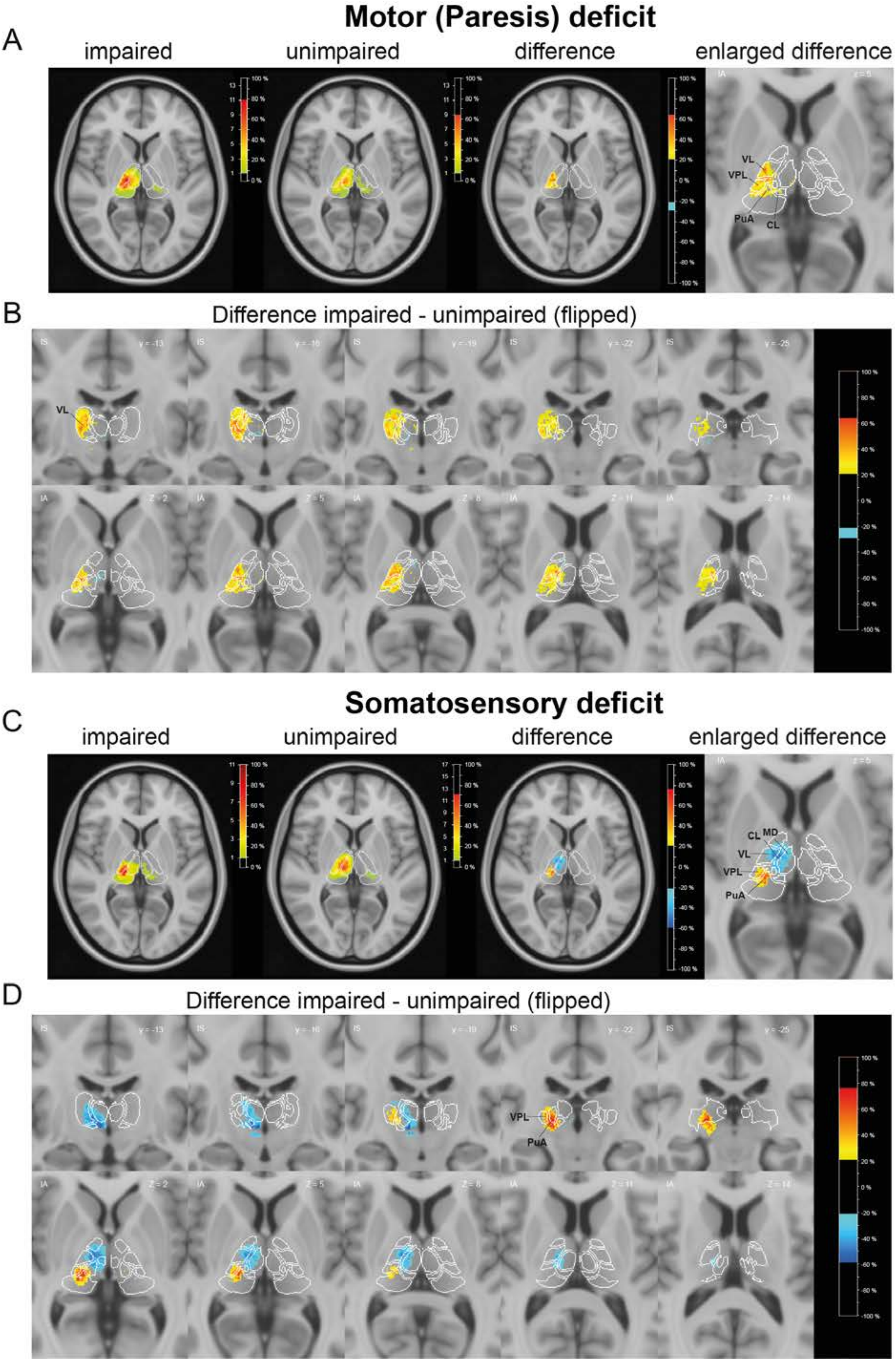

Motor (Paresis) deficit
A
impaired
unimpaired
difference
enlarged difference
VL
VPL
PuA
CL
B
Difference impaired - unimpaired (flipped)
VL
Somatosensory deficit
C
impaired
unimpaired
difference
enlarged difference
CL
MD
VL
VPL
PuA
D
Difference impaired - unimpaired (flipped)
VPL
PuA

**Supplementary Figure 6 Control subtraction for patients with motor and somatosensory deficits (npatients = 28). (A)** Lesion overlap of the patients with ($n_{impaired}$ = 14, $n_{unimpaired}$ = 14) and without an upper limb paresis (sinking of the contralesional arm while holding). **(B)** Percentage of overlapping lesions of the paresis patients after subtraction of the patients without paresis. *Upper panel*: coronal, *lower panel:* axial slices. The colorbar specifies the percentage of patients with lesions in each voxel, with blue indicating the fewest (dark-light blue), and the hot colors (orange-red) a higher number of patients with the deficit. **(C)** Lesion overlap of the patients with ($n_{impaired}$ = 11, $n_{unimpaired}$ = 17) and without a contralesional somatosensory deficit. **(D)** Percentage of overlapping lesions of the patients with somatosensory symptoms after subtraction of the patients without a somatosensory deficit. Coronal (upper panel) and axial slices (lower panel) are shown. The colorbar specifies the percentage of patients with lesions in each voxel, with blue indicating the fewest, and the hot colors (orange-red) a higher number of patients with the deficit. White outlines in (A-D) represent the borders of thalamic nuclei as defined in the digital version of the Morel atlas normalized to the 0.5 mm MNI152 T1 template. Colored schemes of nuclei outlines are depicted in **Supplementary Fig. 5.** Abbreviations: **VL**, Ventral lateral nucleus; **VPL**, ventral posterior lateral nucleus; **MD**, Mediodorsal nucleus; **CL**, Central lateral nucleus; **PuA**, Anterior pulvinar.

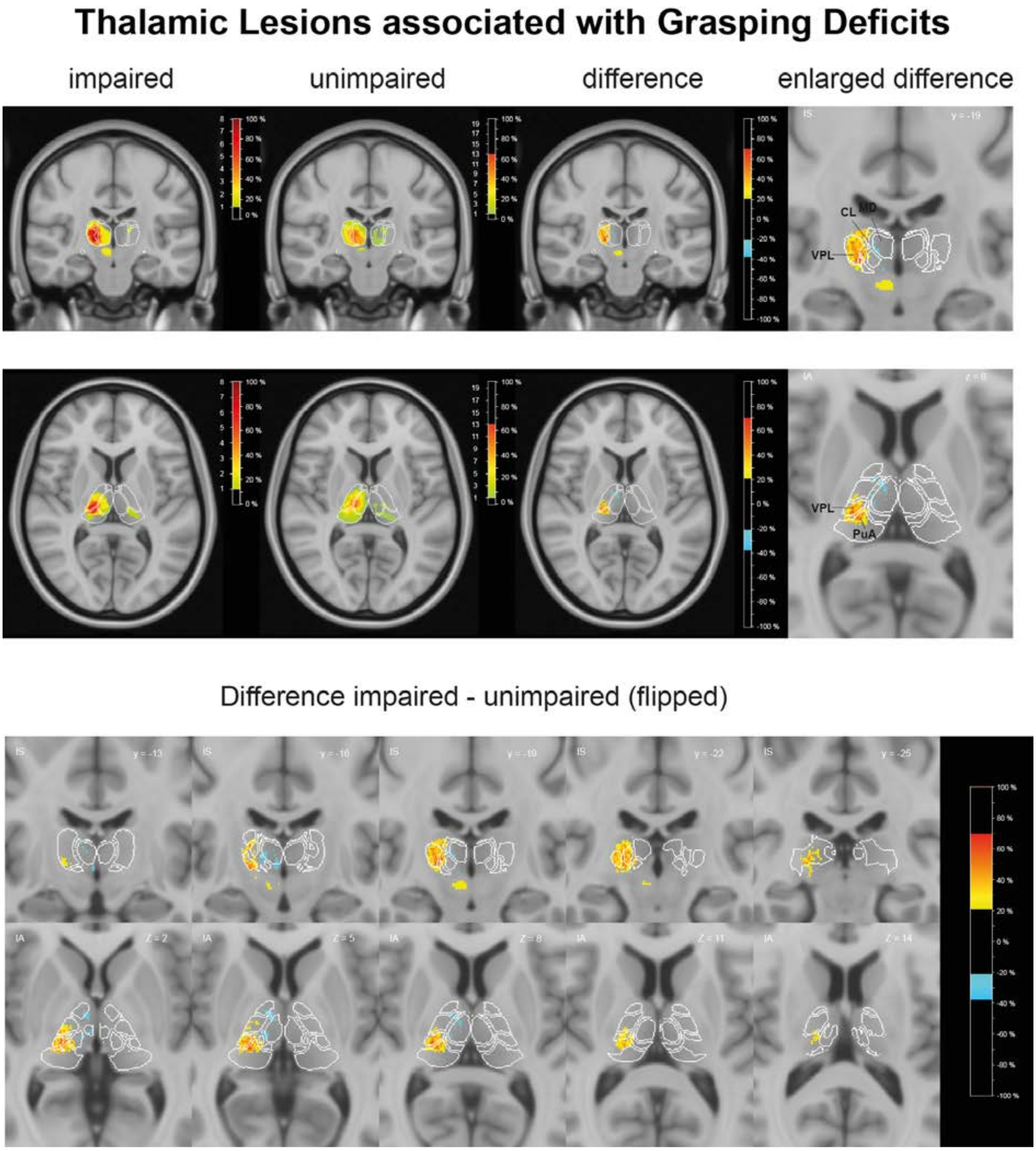


**Supplementary Figure 7 Lesion subtraction plots of the patients with and without grasping errors in the SMARTIES task**. (flipped to the right hemisphere) **(upper row)** Lesion overlap of the patients with (($n_{impaired}$ = 7) and without ($n_{unimpaired}$ = 21) z-scored grasping errors with the contralesional hand. *Upper panel*: coronal, *lower panel:* axial slices. Inset shows the enlarged lesion subtraction map. **(lower row)** Percentage of overlapping lesions of the patients with grasping deficits after subtraction of the patients with normal performance in the grasping task with more slices. The colorbar specifies the percentage of patients with lesions in each voxel, with dark-light blue indicating the fewest, and the hot colors (orange-red) a higher number of patients with a grasping deficit. White outlines represent the thalamic nuclei borders as defined in the digital version of the Morel atlas normalized to the 0.5mm MNI152 T1 template. Abbreviations: **VPL**, ventral posterior lateral nucleus; **MD**, Mediodorsal nucleus; **CL**, Central lateral nucleus; **PuA**, Anterior pulvinar.

### Supplementary Tables

**Supplementary Table 1 Distribution of lesion sites of each patient**

| | Anterior | | | | Medial | | | | | | | | | Posterior | | | | | | | | | | Lateral | | | |
|---|---|---|---|---|---|---|---|---|---|---|---|---|---|---|---|---|---|---|---|---|---|---|---|---|---|---|---|
| **Patient** | **AD** | **AM** | **AV** | **LD** | **MD** | **CM** | **Pf** | **sPf** | **CL** | **CeM** | **Pv** | **MV** | **Hb** | **PuA** | **PuI** | **PuL** | **PuM** | **LP** | **Li** | **Po** | **SG** | **LGN** | **MGN** | **VA** | **VL** | **VM** | **VP** |
| N045L | ++ | | + | +++ | +++ | +++ | +++ | + | +++ | + | | | +++ | +++ | | ++ | +++ | +++ | +++ | +++ | +++ | | + | | ++ | | +++ |
| N049L | | | | | | ++ | + | | | | | | | ++ | | | + | | + | +++ | + | | + | | | | + |
| N054L | | | + | | ++ | + | + | + | ++ | ++ | + | + | + | + | | | | + | | | | | | + | ++ | ++ | + |
| N056L | | | + | + | + | + | | | + | | | | | | | | | | | | | | | | + | | + |
| N056R | | | | | | | | | | | | | | | | | | | | | | | | + | | | |
| N058L | | | | | ++ | +++ | +++ | +++ | + | + | | + | ++ | + | | | | | + | | | | | | + | ++ | + |
| N063L | | | | | | ++ | | | + | | | | | +++ | + | + | + | ++ | | ++ | + | + | ++ | | + | + | +++ |
| N063R | | | | | | + | | | + | | | | | ++ | | + | + | + | | | | | | | | | + |
| N065R | | | | | + | + | | | + | | | | | +++ | | + | + | + | | + | | | + | | + | | ++ |
| N069L | | + | + | | ++ | + | + | +++ | + | +++ | | +++ | | + | | | | | | | | | | ++ | ++ | +++ | + |
| N070R | | | | | + | + | | | + | | | | | +++ | | + | + | ++ | + | ++ | + | | + | | + | | ++ |
| N071R | | | | | + | + | ++ | ++ | | | | | + | | | | | | + | | | | | | | | |
| N072L | | | | | | + | | | | | | | | ++ | | | + | | | +++ | ++ | | ++ | | | | + |
| N074L | | | | + | + | ++ | | | ++ | | | | | +++ | | ++ | + | +++ | | + | | | + | | ++ | + | +++ |
| N075L | | | | | + | + | | | + | | | | | ++ | | | + | + | + | ++ | + | | | | | | + |
| N075R | | | | | + | ++ | | | + | | | | | +++ | | | | | | + | | | | | + | + | ++ |
| N077L | | | | | | | | | | | | | | ++ | | + | + | + | | + | | | + | | | | ++ |
| N078R | | + | | | + | + | +++ | ++ | + | +++ | ++ | +++ | + | | | | | | + | | | | | + | + | + | |
| N080R | | | | | | ++ | | | + | | | | | +++ | | | + | + | | + | | | + | | + | + | +++ |
| N084L | | | | | + | ++ | + | +++ | + | + | | | | + | | | | | | | | | | | + | + | + |
| N086R | | | | | | | | | + | | | | | | | | | + | | | | | | + | ++ | | + |
| N087L | | | | | + | + | ++ | +++ | + | ++ | | +++ | | | | | | | | | | | | + | + | +++ | + |
| N089R | | | | | | + | | | + | | | | | +++ | | + | + | + | | + | | | + | | | | ++ |
| N090L | | | | | | | | | | | | + | | | | | | | | | | | | | | | |
| N090R | | | | | ++ | +++ | +++ | +++ | ++ | +++ | + | +++ | + | + | | | | | + | + | | | | ++ | ++ | +++ | + |
| N093R | | | | | ++ | +++ | +++ | +++ | + | + | | + | + | + | | | | | + | | | | | + | + | +++ | + |
| N095R | | | | + | + | + | | | + | | | | | ++ | | + | + | +++ | | + | | | + | | ++ | + | +++ |
| N096L | | | | + | ++ | +++ | +++ | ++ | ++ | + | | + | + | +++ | | + | +++ | + | +++ | +++ | +++ | | + | | | | + |
| N096R | | | | | + | ++ | +++ | +++ | + | + | | | + | | | | + | | ++ | | + | | | | | | + |
| N097R | | | | | | + | + | | | | | | | + | | | | | + | ++ | ++ | | + | | | | + |
| N098R | | | | | + | ++ | +++ | +++ | + | + | | + | + | | | | | | + | | | | | | + | ++ | + |
| N099L | | | | | | + | + | | + | | | | | + | | + | + | + | + | ++ | ++ | | + | | | | |
| N099R | | | | | | + | | | | | | | | +++ | | | + | + | + | +++ | ++ | | + | | + | + | ++ |
| N106L | | | | | ++ | +++ | +++ | +++ | ++ | + | | + | +++ | +++ | | | + | + | +++ | ++ | + | | | + | ++ | +++ | +++ |

This table delineates the thalamic nuclei impacted by the lesion, classified according to the Morel's neuroanatomical Atlas (2007). The extent of the lesion is presented for each nucleus, denoted as follows: “+” indicates less than 1/3 volume affected, “++” signifies between 1/3 and 2/3 volume affected, and “+++” denotes more than 2/3 of the nucleus volume affected. Abbreviations: **AD**, Anterior dorsal nucleus; **AM**, Anterior medial nucleus; **AV**, Anterior ventral nucleus; **LD**, Lateral dorsal nucleus; **MD**, Mediodorsal nucleus, **CM**, Centromedian nucleus; **Pf**, Parafascicular nucleus; **sPf**, Subparafascicular nucleus; **CL**, Central lateral nucleus; **CeM**, Central medial nucleus; **Pv**, Paraventricular nucleus; **MV**, Medioventral nucleus; **Hb**, Habenular nucleus; **PuA**, Anterior pulvinar; **PuI**, Inferior pulvinar; **PuL**, Lateral pulvinar; **PuM**, Medial pulvinar; **LP**, Lateral posterior nucleus; **Li**, Limitans nucleus; **Po**, Posterior nucleus; **SG**, Suprageniculate nucleus; **LGN**, Lateral geniculate nucleus; **MGN**, Medial geniculate nucleus; **VA**, Ventral anterior nucleus; **VL**, Ventral lateral nucleus; **VM**, Ventral medial nucleus; **VP**, Ventral posterior nucleus. Thalamic nuclei impacted by the lesion were classified according to the Morel's neuroanatomical Atlas [10,11].

**Supplementary Table 2  Frequency of thalamic lesion sites**

| Number of Patients | Patients in % | Nucleus |
|---|---|---|
| 23 | 82.14 | CM |
| 20 | 71.42 | VP |
| 18 | 64.28 | PuA |
| 15 | 53.57 | VL |
| 14 | 50 | CL |
| 13 | 46.42 | Pf |
| 13 | 46.42 | Po |
| 13 | 46.42 | sPf |
| 11 | 39.28 | LP |
| 11 | 39.28 | MD |
| 11 | 39.28 | VM |
| 10 | 35.71 | CeM |
| 8 | 28.57 | Hb |
| 8 | 28.57 | Li |
| 8 | 28.57 | MGN |
| 8 | 28.57 | SG |
| 7 | 25 | MV |
| 6 | 21.42 | VA |
| 4 | 14.28 | PuL |
| 3 | 10.71 | PuM |
| 2 | 7.14 | LD |
| 2 | 7.14 | Pv |
| 1 | 3.57 | AD |
| 1 | 3.57 | AM |
| 0 | 0 | AV |
| 0 | 0 | LGN |
| 0 | 0 | PuI |

Frequency of lesions is plotted for the N = 28 patients. In case of bilateral lesions the largest lesion site was counted. The minimum lesion size was set to 10% of the nucleus. Abbreviations: **AD**, Anterior dorsal nucleus; **AM**, Anterior medial nucleus; **AV**, Anterior ventral nucleus; **LD**, Lateral dorsal nucleus; **MD**, Mediodorsal nucleus, **CM**, Centromedian nucleus; **Pf**, Parafascicular nucleus; **sPf**, Subparafascicular nucleus; **CL**, Central lateral nucleus; **CeM**, Central medial nucleus; **Pv**, Paraventricular nucleus; **MV**, Medioventral nucleus; **Hb**, Habenular nucleus; **PuA**, Anterior pulvinar; **PuI**, Inferior pulvinar; **PuL**, Lateral pulvinar; **PuM**, Medial pulvinar; **LP**, Lateral posterior nucleus; **Li**, Limitans nucleus; **Po**, Posterior nucleus; **SG**, Suprageniculate nucleus; **LGN**, Lateral geniculate nucleus; **MGN**, Medial geniculate nucleus; **VA**, Ventral anterior nucleus; **VL**, Ventral lateral nucleus; **VM**, Ventral medial nucleus; **VP**, Ventral posterior nucleus.

**Supplementary Table 3 Reach error scores and optic ataxia scores for patient groups and healthy controls**

| Variable name | Thalamus Patients | Healthy Controls |
|---|---|---|
| AT_CH_CS_Fov | 0.9 [0.0 17.0] | 0.0 [0.0 4.7] |
| AT_CH_CS_nonFov | 3.4 [0.0 20.5] | 1.2 [0.0 12.1] |
| AT_IH_IS_Fov | 0.9 [0.0 5.7] | 0.0 [0.0 5.0] |
| AT_IH_IS_nonFov | 3.4 [0.0 18.0] | 1.2 [0.0 10.0] |
| AT_CH_IS_Fov | 0.0 [0.0 16.2] | 0.0 [0.0 3.8] |
| AT_CH_IS_nonFov | 0.0 [0.0 16.2] | 0.0 [0.0 3.8] |
| AT_IH_CS_Fov | 0.0 [0.0 16.2] | 0.0 [0.0 3.8] |
| AT_IH_CS_nonFov | 1.0 [0.0 16.2] | 0.0 [0.0 3.8] |
| OA_Score_CH_CS | 2.4 [-2.0 13.6] | 0.5 [-4.7 10.4] |
| OA_Score_IH_IS | 0.7 [-1.4 15.9] | 0.0 [-3.1 7.6] |
| OA_AtaxiaScore_CH_IS | 0.0 [-12.2 19.4] | 0.0 [-2.7 5.9] |
| OA_AtaxiaScore_IH_CS | 0.0 [-2.3 7.2] | 0.0 [-2.9 7.6] |
| AT_PercError_CH_CS_Fov_String_MED | 3.0 [0.0 55.0] | 0.0 [0.0 15.6] |
| AT_PercError_CH_CS_nonFov_String_MED | 10.0 [0.0 69.0] | 0.0 [0.0 25.0] |
| AT_PercError_IH_IS_Fov_String_MED | 0.0 [0.0 16.7] | 2.5 [0.0 25.0] |
| AT_PercError_IH_IS_nonFov_String_MED | 3.7 [0.0 39.3] | 0.0 [0.0 25.0] |
| AT_PercError_CH_IS_Fov_String_MED | 0.0 [0.0 42.9] | 0.0 [0.0 11.5] |
| AT_PercError_CH_IS_nonFov_String_MED | 0.0 [0.0 54.5] | 0.0 [0.0 13.6] |
| AT_PercError_IH_CS_Fov_String_MED | 0.0 [0.0 42.9] | 0.0 [0.0 11.5] |
| AT_PercError_IH_CS_nonFov_String_MED | 2.1 [0.0 20.0] | 0.0 [0.0 16.7] |
| AT_UnCorPercError_CH_CS_Fov_String_MED | 0.0 [0.0 2.5] | 0.0 [0.0 4.5] |
| AT_UnCorPercError_CH_CS_nonFov_String_MED | 0.0 [0.0 14.3] | 0.0 [0.0 7.1] |
| AT_UnCorPercError_IH_IS_Fov_String_MED | 0.0 [0.0 3.6] | 0.0 [0.0 5.9] |
| AT_UnCorPercError_IH_IS_nonFov_String_MED | 0.0 [0.0 13.3] | 0.0 [0.0 10.7] |
| AT_UnCorPercError_CH_IS_Fov_String_MED | 0.0 [0.0 19.2] | 0.0 [0.0 3.8] |
| AT_UnCorPercError_CH_IS_nonFov_String_MED | 0.0 [0.0 18.2] | 0.0 [0.0 8.3] |
| AT_UnCorPercError_IH_CS_Fov_String_MED | 0.0 [0.0 19.2] | 0.0 [0.0 3.8] |
| AT_UnCorPercError_IH_CS_nonFov_String_MED | 0.0 [0.0 8.3] | 0.0 [0.0 6.7] |

Shown is the median and the range. AT_* signifies the reach error scores. Same hand and space conditions ('congruent'): contralesional hand and space: AT_CH_CS; ipsilesional hand and space: AT_IH_IS. Differing hand and space ('incongruent'): ipsilesional hand and contralesional space: AT_IH_CS; contralesional hand and ipsilesional space: AT_CH_IS. The _fov indicates that subjects looked at the pen while reaching and _nonFov indicates the peripheral conditions, when subjects fixated the camera. OA_ stands for the optic ataxia score as a function of hand and space. Percentage of corrected and uncorrected reach errors: AT_PercError_* includes the 'fluent' (score: 0) and 'corrected' (slow, insecure but corrected during reach, score: 1-2); AT_UnCorPercError_ ('uncorrected') includes the scores 3-5, i.e. when the hand stopped at the wrong position and was corrected only in subsequent reaches.

**Supplementary Table 4 Median and range of reach-grasp errors with small objects ("smarties")**

| | Contra hand, contra space | Ipsi hand, Ipsi space | Contra hand, ipsi space | Ipsi hand, contra space |
|---|---|---|---|---|
| **Thalamus Patients** | 1.9 [0.0, 22.0] | 2.2 [0.0, 17.5] | 2.5 [0.0, 23.3] | 0.0 [0.0, 16.3] |
| **with left hemisphere lesion** | 1.2 [0.0, 11.7] | 2.2 [0.0, 11.3] | 1.8 [0.0, 23.3] | 0.6 [0.0, 7.5] |
| **with right hemisphere lesion** | 3.1 [0.0, 22.0] | 1.9 [0.0, 17.5] | 2.5 [0.0, 17.0] | 0.0 [0.0, 16.3] |
| **Healthy Controls** | 0.0 [0.0, 8.8] | 0.0 [0.0, 7.0] | 0.0 [0.0, 3.0] | 0.0 [0.0, 5.0] |

### Supplementary References


1. Fels M, Geissner E. Neglect-Test (NET): ein Verfahren zur Erfassung visueller Neglectphänomene; Handanweisung; deutsche überarbeitete Adaption des Behavioural Inattention Test (Wilson, Cockburn & Halligan, 1987).

2. Bickerton WL, Samson D, Williamson J, Humphreys GW. Separating forms of neglect using the Apples Test: validation and functional prediction in chronic and acute stroke. *Neuropsychology*. 2011;25(5):567-580.

3. Rorden C, Karnath HO. A simple measure of neglect severity. *Neuropsychologia*. 2010;48(9):2758-2763.

4. Vaes N, Lafosse C, Nys G, et al. Capturing peripersonal spatial neglect: an electronic method to quantify visuospatial processes. *Behav Res Methods*. 2015;47(1):27-44.

5. Posner MI, Snyder CR, Davidson BJ. Attention and the detection of signals. *J Exp Psychol*. 1980;109(2):160-174.

6. Rengachary J, He BJ, Shulman GL, Corbetta M. A behavioral analysis of spatial neglect and its recovery after stroke. *Front Hum Neurosci*. 2011;5:29.

7. Rengachary J, d'Avossa G, Sapir A, Shulman GL, Corbetta M. Is the posner reaction time test more accurate than clinical tests in detecting left neglect in acute and chronic stroke? *Arch Phys Med Rehabil*. 2009;90(12):2081-2088.

8. Wilke M, Turchi J, Smith K, Mishkin M, Leopold DA. Pulvinar Inactivation Disrupts Selection of Movement Plans. *Journal of Neuroscience*. 2010;30(25):8650-8659.

9. Wilke M, Schneider L, Dominguez-Vargas AU, et al. Reach and grasp deficits following damage to the dorsal pulvinar. *Cortex*. 2018;99:135-149.

10. Krauth A, Blanc R, Poveda A, Jeanmonod D, Morel A, Székely G. A mean three-dimensional atlas of the human thalamus: generation from multiple histological data. *Neuroimage*. 2010;49(3):2053-2062.

11. Morel A, Magnin M, Jeanmonod D. Multiarchitectonic and stereotactic atlas of the human thalamus. *J Comp Neurol*. 1997;387(4):588-630.
